# Diffusion-Based Synthetic Pretraining for Root Segmentation in Rhizotron Images

**DOI:** 10.64898/2026.10.30.763050

**Authors:** Fengjiao Lu, Gert Kootstra, Zane K. J. Hartley, Rick van de Zedde, Michael Pound, Viola Willemsen

## Abstract

Rhizotrons allow root system architecture to be studied non-destructively in soil. However, automated analysis of rhizotron images remains difficult: roots have low contrast against soil, soil particles resemble fine roots, and deep learning models need many manually annotated images that are slow and subjective to produce. Synthetic images could provide paired images and annotations at scale, but rhizotron images are largely absent from the data used to train generative models. We built a pipeline combining Low-Rank Adaptation (LoRA), ControlNet and inpainting to generate synthetic rhizotron images with paired root masks. We compared training on synthetic data only, real data only, and synthetic pretraining followed by fine-tuning on real images, using three segmentation models: nnU-Net v2, YOLO26 and Mask2Former. Synthetic-only training reached a Dice score of up to 0.60. Synthetic pretraining followed by fine-tuning on the full real training set (two annotated rhizotron images) reached 0.80, and nnU-Net v2 and Mask2Former reached 0.62 with only two annotated image patches containing visible roots. When real annotations were limited, pretraining also improved segmentation of unseen crops, including lettuce, barley, wheat, maize, tomato, cucumber, French bean and field bean. We also introduced a Soft Dice score that compares predictions with probabilistic consensus maps from seven independent annotators, accounting for annotator disagreement. Models matched average human agreement on low-ambiguity patches but remained below it on high-ambiguity patches. In conclusion, diffusion-generated images can complement real annotations and help scale root segmentation, although their effect on downstream root trait measurements still needs to be validated.

## 1 Introduction

Traditionally, root system architecture (RSA) has been studied using transparent agar plates, which enable direct observation of root growth and facilitate quantitative analysis. Although agar-based systems have greatly advanced the understanding of root development, they do not accurately represent the physical and chemical conditions experienced by roots in natural soils. Consequently, rhizotrons have become an increasingly important platform for root phenotyping because they allow plants to grow in soil while maintaining non-destructive visual access to the root system. This enables the investigation of root phenotypes of previously identified root development mutants and responses to environmental stresses under more realistic conditions, and supports the identification of genes associated with stress adaptation and resource acquisition.

Recent developments in automated imaging systems have enabled rhizotrons to be deployed in high-throughput phenotyping platforms, allowing thousands of root images to be acquired throughout plant development. Such large-scale experiments provide opportunities to investigate natural variation, genotype-environment interactions, and dynamic root growth. However, the number of images far exceeds the capacity of manual analysis. While manual annotation may be feasible for small experiments, it rapidly becomes time-consuming for high-throughput studies, creating an urgent need for automated and scalable image analysis methods.

Root segmentation is a fundamental step in automated root phenotyping because accurate segmentation directly determines the reliability of trait measurements. However, rhizotron images pose greater challenges for image segmentation compared to agar plate images. Roots often exhibit low contrast against the surrounding soil, while soil particles, cracks, and organic debris introduce visually similar structures that can easily be mistaken for roots. In addition, fine lateral roots and root hairs occupy only a few pixels, root systems frequently overlap or occlude one another, and illumination conditions can vary across images. These factors make accurate segmentation considerably more difficult than in conventional agar-based imaging systems.

Early root image analysis relied heavily on manual tracing, which is labour-intensive and difficult to scale to high-throughput phenotyping. To reduce this burden, semi-automated tools were developed to assist with root tracing and trait measurement. SmartRoot [1] represents roots as connected biological structures and allows users to trace and correct root systems, while RhizoTrak [2] provides an open-source framework for annotating roots in time-series minirhizotron images. RhizoVision Explorer [3] is an open-source tool for quantifying root traits such as length, diameter, surface area, and volume from scanned root samples. Measurement accuracy can vary with image resolution, root overlap, and analysis settings. These approaches reduced manual effort but still relied substantially on user intervention, motivating the development of fully automated deep-learning methods for root segmentation.

Deep learning has increasingly enabled automated root segmentation and more efficient root phenotyping. Early CNN-based approaches, such as SegRoot [4], demonstrated the potential of deep learning to segment roots in complex soil backgrounds. Subsequent models, including U-Net [5] and an improved DeepLabv3+ architecture [6], further improved segmentation of roots in heterogeneous backgrounds while preserving fine structures. Deep learning has also been extended beyond pixel-wise segmentation: RootNav 2.0 [7] combines segmentation with root-tip detection to reconstruct root trajectories and branching structures, while RootPainter [8] uses human-in-the-loop correction to reduce annotation effort. These developments highlight the potential of deep learning for automated root segmentation and root phenotyping, while also revealing the continuing need for manually annotated data.

The need for manually annotated data has motivated increasing interest in synthetic data as an alternative source of training images and annotations. A key advantage of synthetic data is that root geometry can be explicitly controlled while ground-truth masks are generated automatically. Early studies therefore focused on constructing biologically plausible root systems using predefined models. Lobet et al. [9], for example, generated thousands of root systems with varied architectures and noise levels, demonstrating the potential of synthetic root geometries for evaluating image-analysis and trait-extraction methods.

Subsequent studies incorporated synthetic root structures into segmentation training. Douarre et al. [10] generated synthetic soil-root CT images and showed that matching simulated soil intensity and texture to real images improved transfer to maize data. Similar approaches were developed for MRI. Uzman et al. [11] combined reconstructed root structures with simulated soil noise, whereas Zhao et al. [12] combined synthetic roots with real pure-soil MRI scans to train super-resolution segmentation models. More recently, Hann et al. [13] combined L-system root models with textured soil in Blender to generate paired micro-CT images and segmentation masks. Collectively, these studies show that effective synthetic-to-real transfer requires not only biologically plausible root structures but also realistic representations of the target imaging environment.

Generative models provide an alternative to explicitly simulating the imaging process by learning the relationship between root structures and image appearance. Thesma and Mohammadpour [14] used Pix2PixHD to generate root images from segmentation maps, while Yu et al. [15, 16] used EnlightenGAN for root-image reconstruction and an improved CycleGAN to generate root images across diverse soil backgrounds. In the latter study, improvements were modest for familiar images but substantially greater under new soil-background conditions, highlighting the potential of synthetic image generation to improve generalisation across imaging conditions.

More recently, diffusion models have provided greater flexibility in controlling both image appearance and spatial structure. Hartley et al. [17], for example, combined Stable Diffusion with LoRA fine-tuning and ControlNet to generate synthetic leaf images while preserving target spatial structures. This combination is particularly relevant to root-image synthesis: LoRA enables adaptation to a specific visual domain, while ControlNet constrains the generated image according to a predefined structure. However, this approach was developed for leaf images, and its applicability to soil-based root imagery has not been tested, where fine roots, complex soil texture, occlusion, and low root-soil contrast pose additional challenges.

Several gaps remain in synthetic root image generation. First, existing evidence for synthetic root segmentation comes mainly from CT and MRI, while RGB-based root studies have focused largely on GAN-based augmentation and image reconstruction. Whether diffusion models can generate realistic RGB rhizotron images suitable for root segmentation therefore remains an open question. This is particularly challenging because rhizotron images contain complex soil texture, illumination variation, occlusion, low root-soil contrast, and species-specific variation. In addition, rhizotron images are poorly represented in the large-scale datasets used to pretrain diffusion models, making effective domain adaptation necessary to generate realistic images while preserving root structure. Second, previous studies have primarily evaluated synthetic data in terms of segmentation accuracy or data augmentation, with limited investigation of whether synthetic data can reduce manual annotation requirements or improve generalisation to unseen plant species. Finally, it remains unclear whether models benefiting from synthetic pretraining can achieve segmentation performance comparable to human annotators when inter-annotator variability is explicitly considered.

To address these gaps, we propose a diffusion-based framework that combines LoRA fine-tuning, ControlNet, and inpainting to generate realistic synthetic rhizotron images (Figure 1). Using the generated dataset, we systematically investigate whether synthetic pretraining can (i) improve segmentation performance, (ii) reduce the amount of manually annotated data required for training, (iii) enhance generalisation to unseen crop species, and (iv) achieve human-level performance using a probabilistic consensus-based evaluation framework.

**Figure 1:**
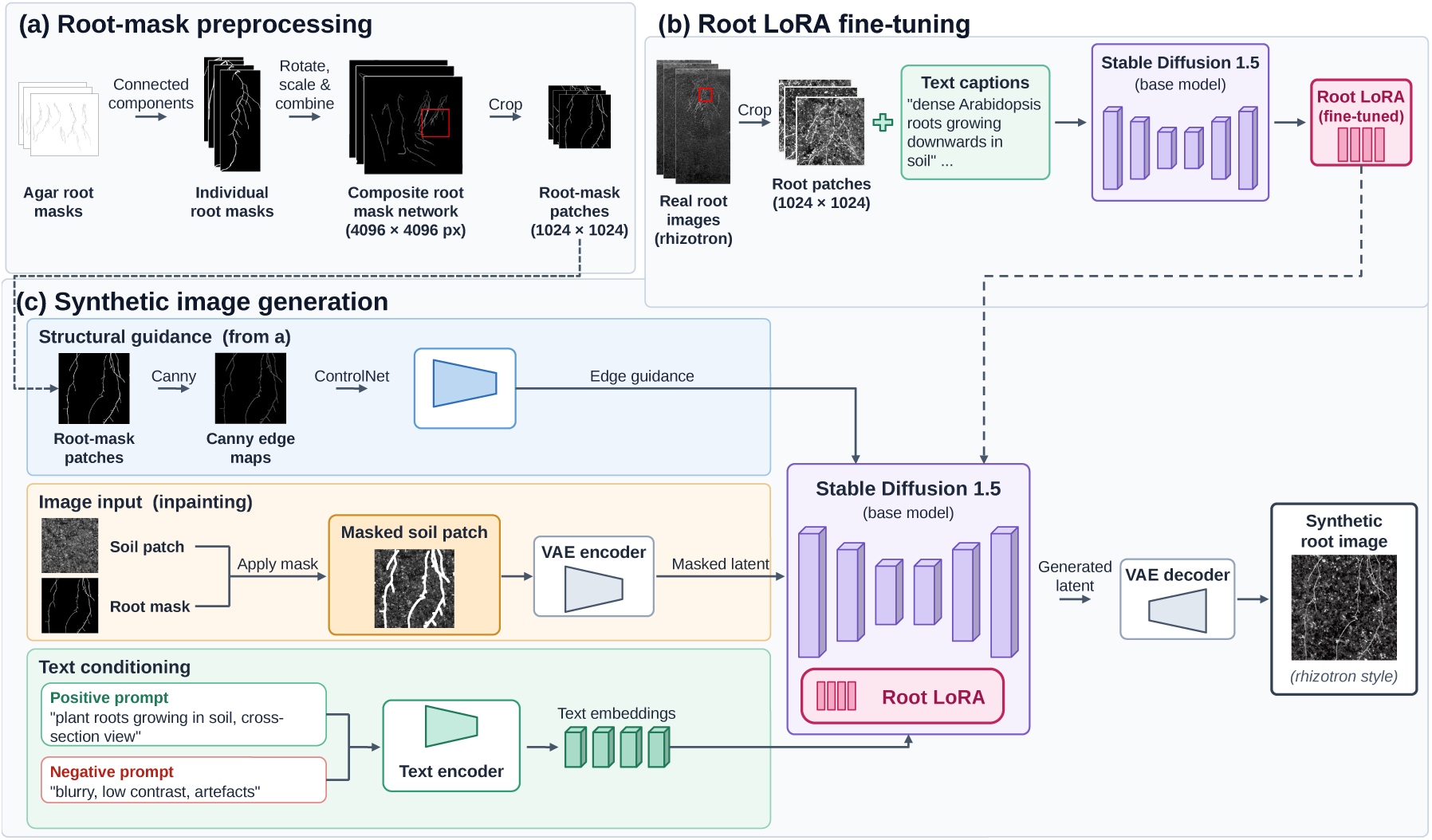
Synthetic rhizotron image-generation pipeline. (a) Root-mask preprocessing: root masks from agar-grown Arabidopsis are rotated, rescaled, combined into networks and cropped into patches. (b) Root LoRA fine-tuning: Stable Diffusion 1.5 is fine-tuned with LoRA on rhizotron root patches and text captions. (c) Synthetic image generation: roots are inpainted into a real soil patch using Stable Diffusion 1.5 with the Root LoRA, guided by the Canny edge map of a root-mask patch through ControlNet. The “Root mask” applied to the soil patch is the inpainting mask (the dilated root-mask patch; see Materials and Methods). The root-mask patch is retained as the segmentation label.

## 2 Materials and Methods

### 2.1 Synthetic Rhizotron Image Generation

We developed a diffusion-based framework for generating synthetic rhizotron images from prescribed root architectures. The framework consists of three components: root-mask preprocessing, Root LoRA fine-tuning, and synthetic image generation (Figure 1). Together, these components combine structurally defined root masks with real soil backgrounds and a diffusion model adapted to rhizotron root appearance, enabling the generation of synthetic images with corresponding segmentation labels.

#### 2.1.1 Root-mask preprocessing

To provide diverse structural guidance for synthetic root generation, we reused an existing set of manually annotated images of Arabidopsis roots grown on agar-plates. These annotations were not generated specifically for the present study; rather, the existing root masks provided a source of root architectures that could be recombined to construct new structural configurations. Individual root systems were extracted using connected-component analysis, randomly rotated and rescaled, and recombined on an empty 4096 *×* 4096-pixel canvas. The number and arrangement of root systems were adjusted to maintain a consistent root pixel density across the generated canvases. The resulting composite root-mask networks were then cropped into 1024 *×* 1024-pixel root-mask patches (Figure 1a).

The root-mask patches were used both to guide image generation and to provide ground-truth masks for segmentation. For generation, each root-mask patch was converted into a Canny edge map, which was provided to ControlNet as structural guidance for the placement and shape of the generated roots. The original root-mask patch was retained unchanged as the corresponding ground-truth root mask for the generated image.

#### 2.1.2 Root LoRA fine-tuning

Images were captured using a WIWAM Rhizotron System equipped with 301 plates. Each Rhizotron plate provides a 4-mm soil layer and an imaging area of 480 mm (height) by 240 mm (width), corresponding to an image resolution of 6400 *×* 13,380 pixels; plates are tilted at 70 degrees to promote root growth along the transparent glass surface. A scanning system with red and blue LED light sources produces two greyscale images, which are combined during post-processing to improve root-soil contrast.

Two types of image patches were prepared from the rhizotron images: root patches and soil patches. Regions of interest (ROIs) containing dense root systems were cropped into 1024 *×* 1024-pixel root patches for Root LoRA fine-tuning, whereas regions without visible root structures were cropped into 1024 *×* 1024-pixel soil patches for use as real soil backgrounds during synthetic image generation.

Stable Diffusion is a diffusion-based generative model that progressively denoises a noisy representation toward samples from a learned image distribution. It performs this process in a lower-dimensional latent space rather than directly in pixel space, enabling efficient image generation [18]. Stable Diffusion 1.5 was used as the base model for Root LoRA fine-tuning and subsequent synthetic image generation.

To adapt the pretrained Stable Diffusion model to the visual characteristics of rhizotron images, we employed Low-Rank Adaptation (LoRA) [19], a parameter-efficient fine-tuning method that keeps the original model weights frozen and trains low-rank adaptation matrices. This allows the pretrained model to be adapted to the specialised rhizotron domain while retaining its general image-generation capabilities, following the plant-specific adaptation strategy demonstrated by Hartley et al. [17].

Root LoRA fine-tuning used the 1024 *×* 1024-pixel root patches obtained from the rhizotron images. Each root patch was paired with a randomly selected text caption describing a dense Arabidopsis root system growing downwards in soil from a side-view rhizotron perspective. The wording of the captions was randomly varied while keeping their biological meaning consistent. Using multiple semantically equivalent captions exposed the model to different textual descriptions of the same visual concept, encouraging the text-conditioning mechanism to associate the captions with the underlying root characteristics rather than with a single fixed wording.

The model was fine-tuned at a resolution of 1024 *×* 1024 pixels. Within Stable Diffusion 1.5, the component responsible for progressively denoising the latent representation during image generation is the U-Net. Only the LoRA adapter weights attached to this denoising component were trained, while the original Stable Diffusion model weights and other pretrained components remained frozen. The LoRA rank and alpha were set to 128 and 64, respectively. Training used the AdamW8bit optimiser with a learning rate of 1 *×* 10*^−^*^4^ and a batch size of 1. Gradients were accumulated over four iterations, giving an effective batch size of 4. A cosine learning-rate scheduler with a 500-step warm-up was used, and the model was trained for 10 epochs using bfloat16 mixed precision. Caption shuffling and a caption dropout rate of 0.1 were applied to introduce additional variation during text-conditioned training. The resulting Root LoRA was used with Stable Diffusion 1.5 during synthetic image generation, together with root-mask patches, soil patches, ControlNet guidance, and text conditioning.

#### 2.1.3 Synthetic image generation

Synthetic rhizotron images were generated using an inpainting pipeline based on Stable Diffusion 1.5 (v1-5-pruned-emaonly). The Root LoRA was applied to the base model to adapt image generation toward the visual characteristics of Arabidopsis roots in rhizotron images.

To control the spatial arrangement of the generated roots, we additionally used ControlNet [20], a neural network architecture that provides spatial conditioning to a pretrained diffusion model. Rather than relying only on text conditioning to determine where objects appear in an image, ControlNet allows structural information, such as edges or segmentation-derived features, to guide the generation process. In this study, root-mask patches were converted into Canny edge maps and provided to ControlNet as structural guidance. This allowed the generated roots to follow the spatial arrangement specified by the root-mask patches while the Root LoRA adapted their visual appearance to the rhizotron domain.

For each synthetic image, a 1024 *×* 1024-pixel soil patch was randomly sampled from the pool of real soil patches to provide the background, and a root-mask patch generated during root-mask preprocessing defined the target root architecture. The root-mask patch was retained unchanged as the segmentation label. For inpainting, the root-mask patch was dilated by 6 pixels and processed with a 60-pixel context padding to form the inpainting mask, which marks the region in which roots are generated; the dilation provides additional context around the root boundaries and reduces edge artefacts.

The inpainting mask was applied to the soil patch, blanking out the pixels to be regenerated, to give the masked soil patch. The masked soil patch was processed by the VAE encoder to obtain the masked latent representation used by the inpainting pipeline. The root-mask patch was also converted into a Canny edge map, which was provided to the ControlNet inpainting model (control-v11p-sd15-inpaint) as structural guidance. Thus, the root-mask patch defined the target root architecture through ControlNet, while the soil patch provided the real soil appearance surrounding the generated roots.

Text conditioning was provided using a positive prompt describing plant roots growing in soil from a cross-section view and a negative prompt to suppress undesirable image characteristics. The positive prompt was “plant roots growing in soil, cross-section view”, while the negative prompt was “blurry, low contrast, artefacts”. The text was encoded into text embeddings and supplied to the diffusion model together with the masked latent representation, Root LoRA, and ControlNet guidance.

Images were generated using the Euler sampler with 20 denoising steps, a classifier-free guidance (CFG) scale of 7.9, and a denoising strength of 1.0. The ControlNet conditioning strength was set to 1.1 to encourage adherence to the root-mask layout. The resulting latent representations were decoded by the VAE decoder to produce the final synthetic root images with rhizotron-style soil backgrounds, yielding paired synthetic images and root masks for subsequent segmentation-model training.

### 2.2 Root Segmentation Training and Evaluation

The generated synthetic images were used to train and pretrain root segmentation models under different training strategies. The models were subsequently evaluated across a series of experiments designed to assess the contribution of synthetic data, annotation efficiency, cross-species generalisation, and agreement with human annotators (Section 2.2.5). An overview of the segmentation training and evaluation framework is shown in Figure 2.

**Figure 2:**
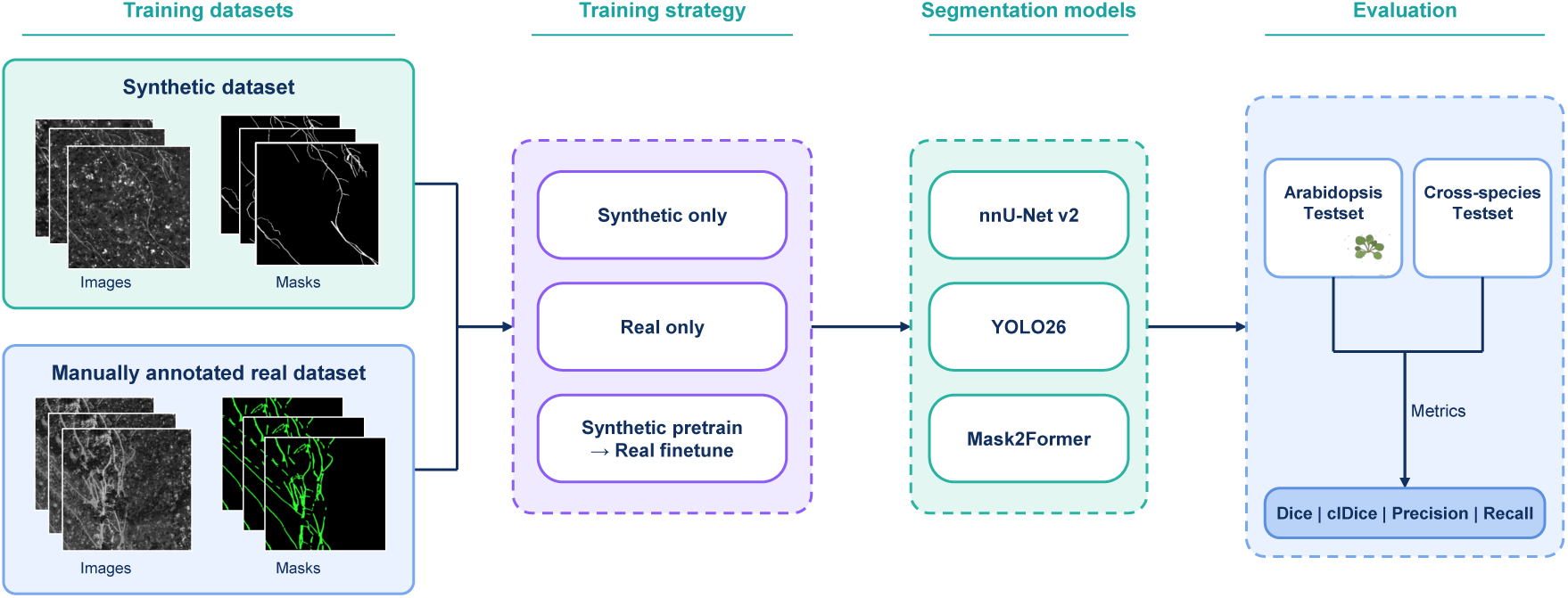
Overview of segmentation training and evaluation in Experiments 1–3. Synthetic and real datasets are used for synthetic-only, real-only, and synthetic-pretraining plus fine-tuning strategies. Models are evaluated on Arabidopsis and cross-species test sets using Dice, clDice, precision and recall. The cross-species test set comprises 30 rhizotron images of eight crops not seen during training (lettuce, barley, wheat, maize, tomato, cucumber, French bean and field bean), together with Arabidopsis as a reference. Human-consensus evaluation is shown in Figure 3.

#### 2.2.1 Training, validation, and test datasets Training and validation datasets

- **Real Arabidopsis dataset:** Two full-resolution (6400 *×* 13,380 pixels) rhizotron images were cropped into 108 patches. These patches were semi-automatically annotated and formed the real-image training dataset.
- **Synthetic dataset:** Approximately 1,000 root-mask patches were used to generate the synthetic rhizotron dataset. To increase the diversity of the generated images, each root-mask patch was combined with multiple randomly selected real soil patches and conditioned using a text prompt describing roots growing in soil (“plant roots growing in soil, cross-section view”). This procedure generated six unique 1024 *×* 1024 pixel synthetic images for each root-mask patch. From the resulting image pool, 3,758 images were selected by taking one image per root-mask patch in turn, so that all root architectures were represented. These images formed the synthetic dataset; for each segmentation model, 80% (3,006 images) were randomly assigned to training and 20% (752 images) to validation.

#### Test datasets

- **Arabidopsis test dataset:** To evaluate baseline performance, 15 full-resolution Arabidopsis rhizotron images were cropped into 810 patches. These were manually annotated to provide pixel-level ground truth.
- **Cross-species test dataset:** To assess model generalisation, an additional dataset of 30 rhizotron images (1620 patches) was constructed from eight crop species (lettuce, barley, wheat, maize, tomato, cucumber, French bean, and field bean) and Arabidopsis, which was included as a reference for the species used in training. These images were cropped into patches and semi-automatically annotated in the V7 Darwin platform.

#### 2.2.2 Segmentation models

For rhizotron root image analysis, semantic segmentation was used to separate root structures from the surrounding soil background. In this study, nnU-Net v2 [21], YOLO26 [22], and Mask2Former [23] were selected to evaluate how synthetic image datasets perform across different segmentation architectures. These three approaches represent different deep learning paradigms and were chosen to test whether the synthetic dataset could support robust learning under different model designs.

##### nnU-Net v2

nnU-Net is a self-configuring segmentation framework that automatically adapts its preprocessing, network architecture, and training strategy to a given dataset. Owing to its robustness and minimal manual configuration, it has become a widely adopted baseline for biomedical image segmentation. Because plant roots exhibit thin, elongated structures similar to blood vessels and other tubular anatomical structures, nnU-Net provides a strong reference framework for rhizotron root segmentation.

##### YOLO26

YOLO26 represents a one-stage segmentation framework designed for efficient end-to-end prediction. Compared with encoder-decoder segmentation networks, YOLO26 emphasises computational efficiency while maintaining competitive segmentation accuracy. It was included to investigate whether synthetic pretraining also benefits lightweight models suitable for high-throughput rhizotron phenotyping applications.

##### Mask2Former

Mask2Former is a transformer-based segmentation framework that employs masked attention to integrate local image features with long-range contextual information. Its unified architecture supports semantic, instance, and panoptic segmentation within a single framework. Such characteristics are particularly beneficial for rhizotron images, where thin root structures are frequently interrupted by low contrast, soil particles, and complex background textures.

#### 2.2.3 Training protocol

The training strategies shown in Figure 2 were used in Experiments 1–3; Experiment 4 used the separate human-consensus evaluation workflow in Figure 3. Three training strategies were used to assess the contribution of synthetic data to root segmentation. First, models were trained exclusively on the synthetic dataset (synthetic-only). Second, models were trained exclusively on the real annotated rhizotron dataset (real-only), providing the baseline condition. Third, models were pretrained on the synthetic dataset and subsequently fine-tuned on the real dataset (synthetic pretraining followed by fine-tuning, abbreviated PT+FT; “Synthetic pretrain *→* Real finetune” in Figure 2). These strategies were applied to nnU-Net v2, YOLO26, and Mask2Former.

**Figure 3:**
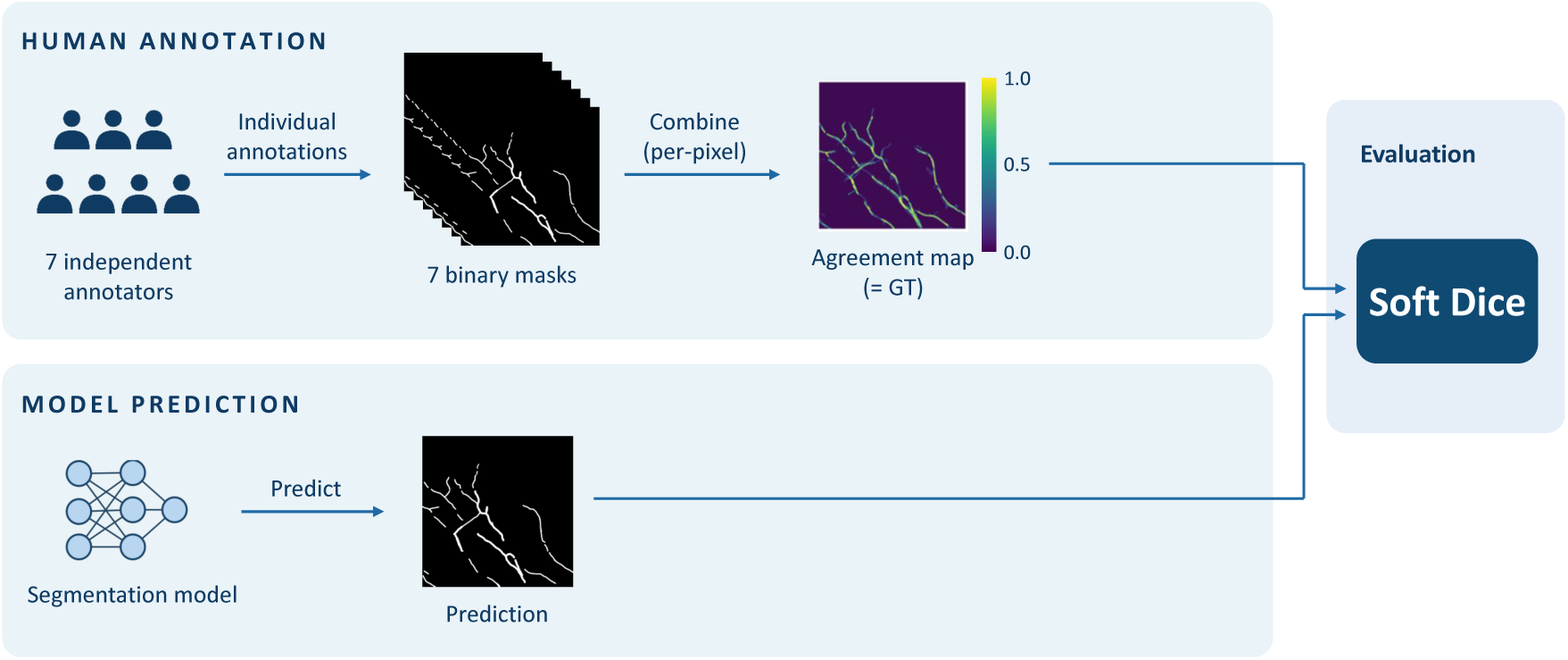
Human-level evaluation in Experiment 4. Seven independent annotations are combined into a per-pixel probabilistic consensus map (labelled “Agreement map” in the figure), which serves as the reference (GT) and is compared with model predictions using Soft Dice. Each annotator is excluded from their own consensus reference.

Training followed the standard configurations of the respective segmentation frameworks, with model-specific settings selected to accommodate the 1024 *×* 1024-pixel input images. nnU-Net v2 automatically configured a 2D PlainConv U-Net with a batch size of 3 and 1024*×*1024-pixel patches, and the model was trained for the default 1000 epochs. YOLO26 used the YOLO26s segmentation model, initialised from COCO-pretrained weights, with a batch size of 16, an input resolution of 1024 *×* 1024 pixels, and 1000 training epochs; for fine-tuning, YOLO26 was trained for 500 epochs with an initial learning rate of 0.001 and a cosine learning-rate schedule. Mask2Former used a ResNet-101 backbone with a batch size of 2, a learning rate of 1*×*10*^−^*^4^, and 80,000 training iterations. The learning rate was reduced at 60,000 and 72,000 iterations, with 10 warm-up iterations. No additional hyperparameter optimisation was performed for comparison between training strategies. At inference, predicted root probabilities were binarised at a threshold of 0.01. Root LoRA training, synthetic image generation and all segmentation experiments were performed on a single NVIDIA RTX A6000 GPU (48 GB).

To investigate whether synthetic pretraining reduces the amount of manually annotated data required for root segmentation, the real training dataset was divided into subsets corresponding to five annotation levels: 10%, 25%, 50%, 75%, and 100%. The complete training dataset consisted of 108 image patches (1024 *×* 1024 pixels), derived from two full-resolution rhizotron images (6400 *×* 13,380 pixels), and was defined as the 100% setting. Of these 108 patches, 35 contained visible roots and 73 contained no visible roots. The patches were ranked according to the number of annotated root pixels, and subsets were constructed to represent 10%, 25%, 50%, 75%, and 100% of the total annotated root pixels. These subsets contained 9, 21, 43, 67, and 108 patches, respectively, including 2, 3, 7, 13, and 35 patches containing visible roots. For each annotation level, model performance was compared between training on the corresponding real-data subset alone (real-only) and synthetic pretraining followed by fine-tuning on the same subset.

#### 2.2.4 Evaluation metrics

##### Root segmentation metrics

Model performance was evaluated using both pixel-wise and topology-aware segmentation metrics. The pixel-wise metrics included the Dice coefficient, precision, and recall, which quantify the agreement between the predicted segmentation and the ground-truth annotation. The Dice coefficient measures the spatial overlap between the predicted and reference masks, whereas precision and recall quantify the accuracy and completeness of the predicted root pixels, respectively. The Dice coefficient was computed as (Equation 1):

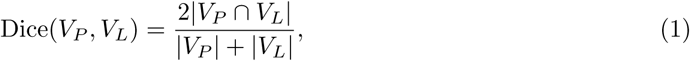

where *V_P_*and *V_L_* denote the predicted and ground-truth foreground pixels, respectively.

Although these metrics provide a comprehensive evaluation of pixel-level segmentation accuracy, they can be overly sensitive for rhizotron root images. Root boundaries are often ambiguous because fine root hairs, low image contrast, and soil particles make it difficult to determine whether individual pixels should be labelled as root or background. Consequently, manual annotations may vary among annotators, particularly along root boundaries, leading to noticeable differences in pixel-wise metrics despite only minor visual differences in the segmentation.

To complement the pixel-wise evaluation, we also employed the centreline Dice (clDice) metric [24], which measures the topological consistency between the predicted and reference segmentations. Unlike conventional overlap-based metrics, clDice evaluates whether the connectivity of thin, tubular structures is preserved by comparing each segmentation mask with the skeleton of the other mask. The topology precision (*T*_prec_) and topology sensitivity (*T*_sens_) are defined as

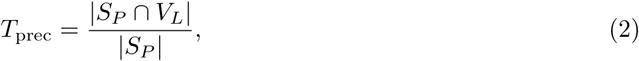

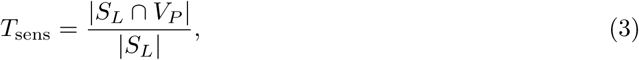

where *S_P_* and *S_L_* denote the skeletons of the predicted and ground-truth masks, respectively. The clDice score is then computed as

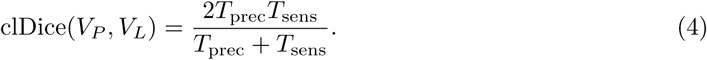

Because plant root systems consist of elongated and highly branched structures, preserving connectivity is often more important than achieving perfect pixel-level agreement. Therefore, clDice provides complementary information to conventional pixel-wise metrics.

##### Human-level evaluation

Manual annotation of rhizotron root images is inherently subjective, particularly along root boundaries and in regions containing fine root hairs. Different annotators may reasonably disagree on whether a given pixel should be labelled as root or background, resulting in variability among reference annotations. In this study, only a single manually annotated mask was used to train the segmentation models, whereas annotations from multiple independent annotators were reserved exclusively for evaluating annotation uncertainty and human-level performance. The training masks were generated by the primary annotator who prepared the real training dataset, ensuring a consistent annotation protocol across all training experiments. The human-level evaluation workflow is shown in Figure 3.

##### Soft Dice for uncertainty-aware evaluation

To account explicitly for inter-annotator variability, we introduce a Soft Dice metric that represents the reference annotation as a probabilistic consensus map rather than as a binary mask. This formulation is designed for segmentation tasks in which the location of object boundaries is inherently uncertain and multiple valid annotations may differ at the pixel level. For each pixel *x*, the consensus probability *P* (*x*) is calculated as (Equation 5)

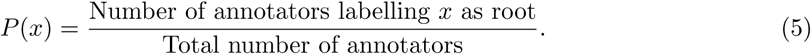

A value of *P* (*x*) = 1 indicates complete agreement that the pixel belongs to the root class, whereas *P* (*x*) = 0 indicates complete agreement that it belongs to the background. Intermediate values represent increasing levels of inter-annotator uncertainty.

In addition, the metric assigns each pixel a confidence weight based on the level of annotator agreement (Equation 6):

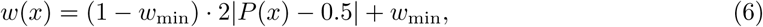

where *w*_min_ = 0.1 specifies the minimum weight assigned to highly uncertain pixels. Pixels for which annotators strongly agree (*P* (*x*) *≈* 0 or *P* (*x*) *≈* 1) therefore receive higher weights, whereas pixels with low agreement receive smaller weights while still contributing to the evaluation. The Soft Dice score is then calculated as (Equation 7)

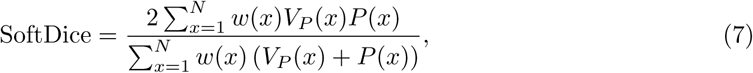

where *V_P_*(*x*) denotes the predicted segmentation value for pixel *x*, and *N* is the total number of pixels in the image.

The metric differs from the conventional Dice coefficient in two respects. First, the reference is represented as a probability derived from multiple independent annotations rather than as a single binary mask. Second, the contribution of each pixel is weighted according to inter-annotator agreement. Consequently, disagreements in regions where annotators themselves exhibit substantial uncertainty have a smaller effect on the final score, while regions with strong human agreement retain greater influence. When evaluating an individual annotator, that annotator is excluded from the consensus map used as the reference (leave-one-out), providing a consistent basis for comparing model and human performance.

#### 2.2.5 Experimental settings

Four experiments were conducted to evaluate the synthetic rhizotron dataset. These experiments assessed (i) the contribution of synthetic data to model training, (ii) annotation efficiency under limited real-data availability, (iii) model generalisation to unseen plant species, and (iv) agreement between model predictions and human annotations.

##### Experiment 1: contribution of synthetic data

This experiment evaluated whether the generated synthetic images provided useful training information for root segmentation. The three training strategies described in Section 2.2.3 were compared using nnU-Net v2, YOLO26, and Mask2Former. Model performance was evaluated on the Arabidopsis Rhozptron test dataset using the segmentation metrics described in Section 2.2.4.

##### Experiment 2: annotation-efficient learning

This experiment investigated whether synthetic pretraining reduces the amount of manually annotated real data required for accurate root segmentation. Models pretrained on the synthetic dataset were fine-tuned using the reduced real-data subsets (100%, 75%, 50%, 25%, and 10%) described in Section 2.2.3 and compared with models trained from only real data using the same subsets.

##### Experiment 3: cross-species generalisation

This experiment evaluated the ability of the trained models to generalise to plant species that were not represented in the training data. All models were trained only on Arabidopsis data (real Arabidopsis Rhizotron images, with or without synthetic pretraining) and subsequently evaluated on the independent cross-species test dataset described in Section 2.2.1, without any additional fine-tuning or domain adaptation. This evaluation assessed the robustness of the models to variations in root morphology, growth patterns, and image characteristics across different plant species. Performance was compared across the two training strategies to determine whether synthetic pretraining improves cross-species generalisation. To quantify the effect, the change in Dice score (ΔDice) was calculated as the difference between the Dice score of the synthetic-pretrained model and that of the corresponding real-only model (Equation 8):

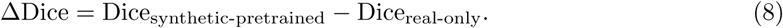

A positive ΔDice indicates an improvement in cross-species segmentation performance following synthetic pretraining, whereas a negative value indicates a decrease.

##### Experiment 4: effect of synthetic pretraining on agreement with human annotations

This experiment evaluated whether synthetic pretraining enabled segmentation models to achieve performance comparable to that of human annotators. Seven independent annotators manually segmented the same set of Arabidopsis rhizotron image patches. The evaluation dataset consisted of two subsets: (i) 10 high-ambiguity image patches containing ambiguous root boundaries and fine root hairs, and (ii) 10 low-ambiguity image patches with clearly distinguishable root structures. The manual annotations were combined to construct the probabilistic consensus maps described in Section 2.2.4, and segmentation performance was evaluated using the Soft Dice metric.

To ensure an unbiased evaluation of human performance, a leave-one-out strategy was adopted. When evaluating an individual annotator, the probabilistic consensus map was generated from the remaining six annotators, excluding the annotation being assessed. For the segmentation models, each predicted mask was compared with each of the seven leave-one-out consensus maps, and the Soft Dice scores were averaged over the seven maps. This protocol enabled a direct comparison between model predictions and human annotations while explicitly accounting for inter-annotator variability.

## 3 Results

### 3.1 Generated Synthetic Images

Figure 4 presents representative synthetic rhizotron images generated using the LoRA–ControlNet inpainting pipeline. The generated images preserve realistic soil texture while producing root systems that closely follow the input root-mask patches. By combining different soil patches with recombined root architectures, the pipeline generated diverse training images exhibiting varying root densities, branching patterns, and spatial arrangements. These synthetic images were subsequently used to evaluate their effectiveness for training root segmentation models.

**Figure 4:**
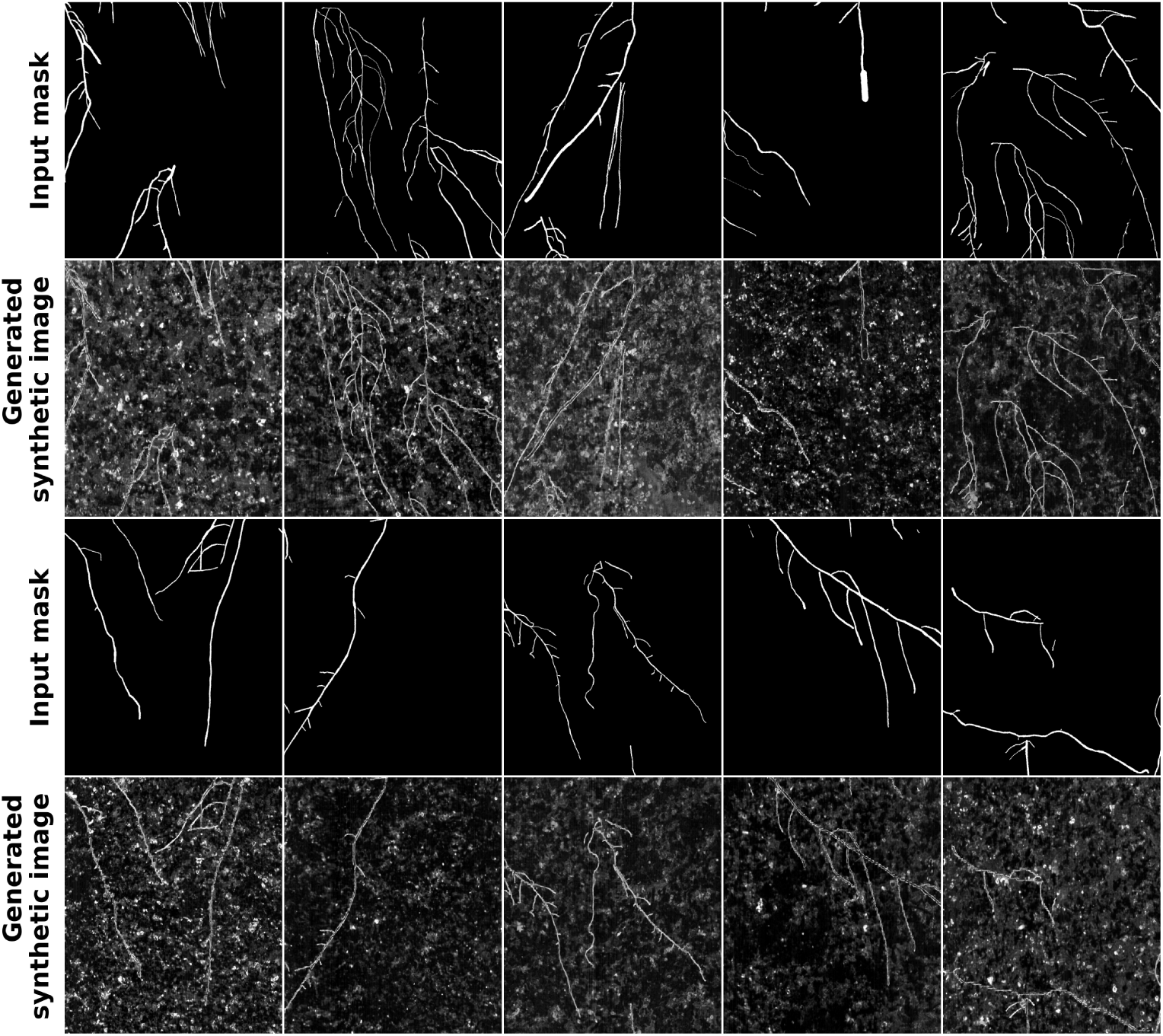
Representative synthetic rhizotron images generated by the proposed pipeline. Rows labelled “Input mask” show root-mask patches; the row below each shows the corresponding generated synthetic image. Images were generated using the Root LoRA, ControlNet root-mask guidance and Stable Diffusion 1.5 inpainting, with real soil patches as backgrounds.

### 3.2 Contribution of Synthetic Data

Experiment 1 compared three training strategies (Figure 2; see Materials and Methods for the experimental design). The quantitative results are summarised in Table 1. Models trained exclusively on synthetic images achieved lower segmentation accuracy than those trained on real images for all three architectures, indicating that the synthetic dataset alone was insufficient to fully replace real annotated data. Nevertheless, the synthetic-only models were able to learn meaningful root representations. Among the three architectures, YOLO26 achieved the highest performance when trained solely on synthetic data, obtaining a Dice score of 0.596 and a clDice score of 0.689, whereas nnU-Net v2 and Mask2Former achieved Dice scores of 0.416 and 0.375, respectively. Complete results, including precision and recall, are given in Supplementary Tables S1–S3.

**Table 1:** Dice and clDice scores for three training strategies on the Arabidopsis test set. Synthetic- only, trained on synthetic data only (no real annotated images); real-only, trained on 100% of the real annotated training dataset (no synthetic data); PT+FT, pretrained on synthetic data and subsequently fine-tuned on 100% of the real annotated training dataset. Bold indicates the highest score within each model–metric pair.

|  | Synthetic-only |  | Real-only |  | PT+FT |  |
|---|---|---|---|---|---|---|
| Model | Dice | clDice | Dice | clDice | Dice | clDice |
| nnU-Net v2 | 0.416 | 0.672 | 0.786 | 0.864 | **0.799** | **0.879** |
| YOLO26 | 0.596 | 0.689 | **0.671** | **0.838** | 0.662 | 0.836 |
| Mask2Former | 0.375 | 0.605 | **0.748** | **0.817** | 0.747 | 0.803 |

When synthetic pretraining was followed by fine-tuning on the real dataset, the largest improvement was observed for nnU-Net v2. Compared with training on real images alone, synthetic pretraining increased the Dice score from 0.786 to 0.799 and the clDice score from 0.864 to 0.879, achieving the best overall performance among all evaluated models. In contrast, synthetic pretraining produced little change for YOLO26 and Mask2Former when the complete real training dataset was available. YOLO26 showed a slight decrease in Dice score (0.671 to 0.662), while Mask2Former achieved nearly identical performance under the two training strategies.

Overall, these results indicate that synthetic pretraining can improve segmentation performance, although its effectiveness depends on the model architecture. Among the evaluated models, nnU-Net v2 benefited most from synthetic pretraining, whereas the advantages for YOLO26 and Mask2Former were limited when sufficient real annotated data were available.

### 3.3 Annotation-Efficient Learning

Figure 5 summarises the Dice and clDice scores obtained under different annotation settings. Synthetic pretraining provided the greatest benefit when only limited annotated real data were available. For nnU-Net v2, the Dice score increased from 0.573 to 0.623 when only 10% of the real training data were used, while the corresponding clDice score increased from 0.643 to 0.703. Similarly, Mask2Former showed a substantial improvement at the 10% annotation level, with the Dice score increasing from 0.460 to 0.624 and the clDice score from 0.527 to 0.691. As the amount of annotated real data increased, the performance gap between the two training strategies gradually decreased, and both approaches achieved comparable performance when the full training dataset was available.

**Figure 5:**
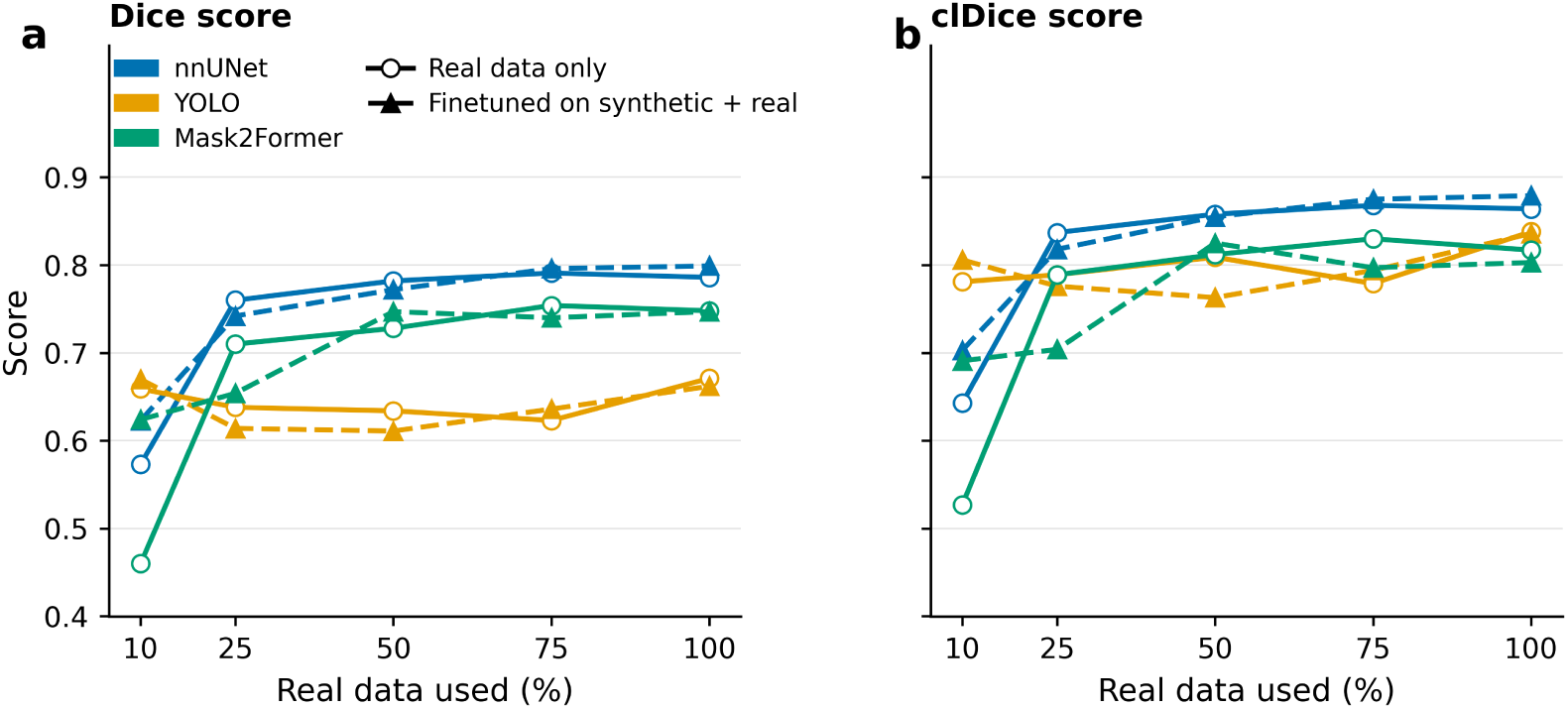
Segmentation performance across real-annotation levels. Open circles (“Real data only”): real-only models trained on the real-data subset; filled triangles (“Finetuned on synthetic + real”): PT+FT models pretrained on synthetic images and fine-tuned on the same subset. The x-axis (“Real data used”) gives the annotation level, defined as the proportion of annotated root pixels in the real training dataset (see Materials and Methods). (a) Dice. (b) clDice. In the legend, nnUNet denotes nnU-Net v2 and YOLO denotes YOLO26.

In contrast, YOLO26 exhibited only minor differences between the two training strategies across all annotation levels. Synthetic pretraining neither improved nor degraded performance, suggesting that its benefit was model dependent.

Overall, synthetic pretraining provided the greatest benefit under annotation-limited conditions, particularly for nnU-Net v2 and Mask2Former.

### 3.4 Cross-Species Generalisation

Average Dice scores on the cross-species test set are summarised in Supplementary Table S4, and the distribution of ΔDice across species is shown in Figure 6. When only 10% of the real training data were available, synthetic pretraining improved segmentation performance for all three models. The largest improvement was observed for Mask2Former, whose average Dice score increased from 0.334 to 0.441 (+0.107). Smaller improvements were observed for YOLO26 (0.599 to 0.626, +0.027) and nnU-Net v2 (0.515 to 0.536, +0.021).

**Figure 6:**
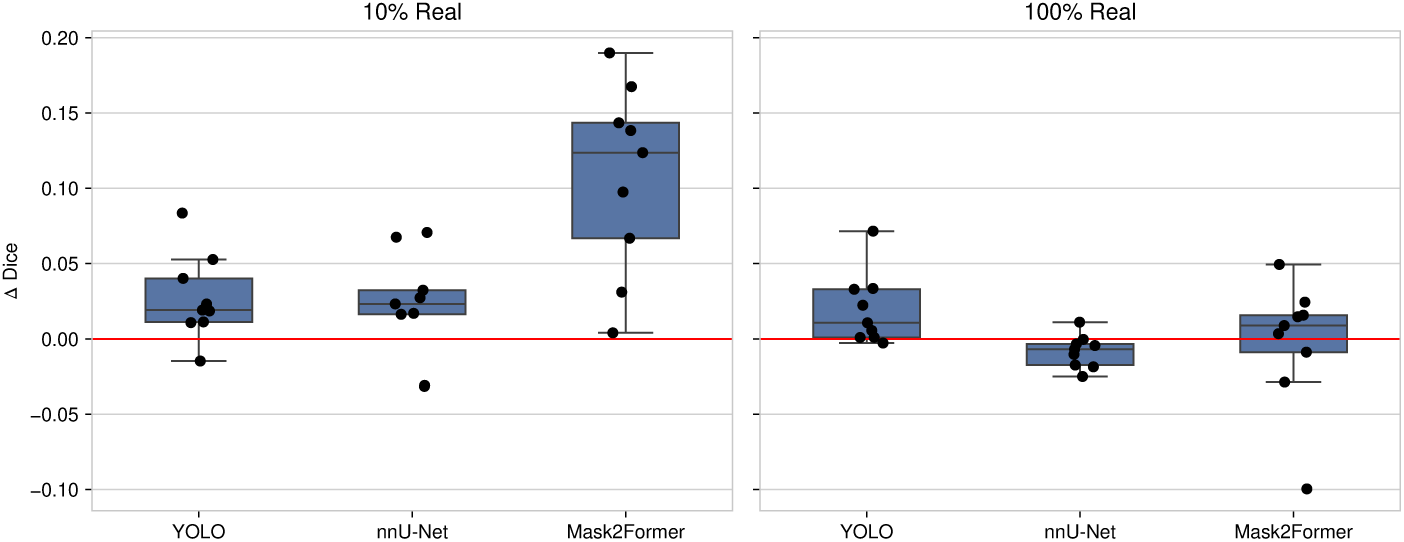
Change in Dice score (ΔDice) for cross-species evaluation. ΔDice is the Dice score of the PT+FT model minus that of the real-only model (Equation 8); each point is one species (Arabidopsis or one of the eight crops). Positive values indicate improvement from synthetic pretraining. “10% Real” and “100% Real” denote the 10% and 100% annotation levels. nnU-Net denotes nnU-Net v2 and YOLO denotes YOLO26.

When the complete real training dataset was used, the effect of synthetic pretraining became considerably smaller. YOLO26 showed a slight improvement (0.648 to 0.668), whereas nnU-Net v2 (0.676 to 0.667) and Mask2Former (0.604 to 0.602) achieved comparable performance with and without synthetic pretraining. The distribution of ΔDice values (Figure 6) further shows that improvements across individual species were substantially larger under the 10% annotation setting than when the complete training dataset was available. Per-species Dice and clDice scores are shown in Supplementary Figs. S1–S2.

The qualitative examples in Figure 7 illustrate the corresponding cross-species predictions for lettuce, cucumber, French bean, maize, wheat, and barley, alongside a prediction for an Arabidopsis test image for reference. The red overlays indicate the root regions predicted by the model trained on Arabidopsis, showing that root structures could be identified across multiple previously unseen crop species without additional fine-tuning or domain adaptation.

**Figure 7:**
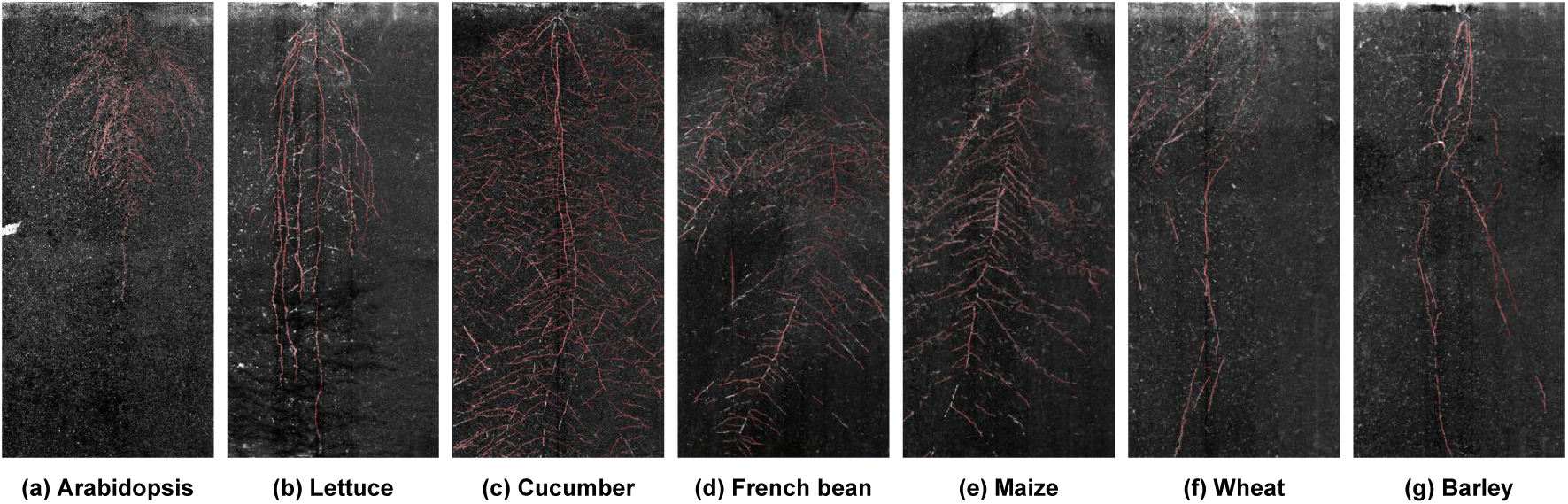
Qualitative segmentation of Arabidopsis and of unseen crops without fine-tuning. Predictions of the Arabidopsis-trained model for (a) Arabidopsis (test image, shown for reference), and for the unseen crops (b) lettuce, (c) cucumber, (d) French bean, (e) maize, (f) wheat and (g) barley, overlaid in red on the original images.

Overall, synthetic pretraining provided the greatest benefit for cross-species segmentation when the amount of real annotated training data was limited. The effect was less pronounced when the complete real training dataset was available. Despite being trained on Arabidopsis, the models were able to generalise to previously unseen crop species without additional fine-tuning or domain adaptation. These findings suggest that synthetic pretraining can provide useful transferable information for cross-species root segmentation, particularly when real annotated data are limited.

### 3.5 Human-Level Evaluation

Experiment 4 compared model predictions with the probabilistic consensus of seven independent annotators using Soft Dice (Figure 3). Figure 8 presents representative high-ambiguity and low-ambiguity image patches together with their corresponding probabilistic consensus maps. Values close to 1 indicate strong agreement that a pixel belongs to the root, whereas values close to 0 indicate agreement that it belongs to the background. Intermediate values indicate increasing dis-agreement among annotators, with values around 0.5 representing the greatest disagreement. The high-ambiguity patches contain more intermediate-valued regions, particularly around fine root hairs and low-contrast root boundaries. In contrast, the low-ambiguity patches are dominated by pixels with values close to 0 or 1, indicating greater agreement among annotators.

**Figure 8:**
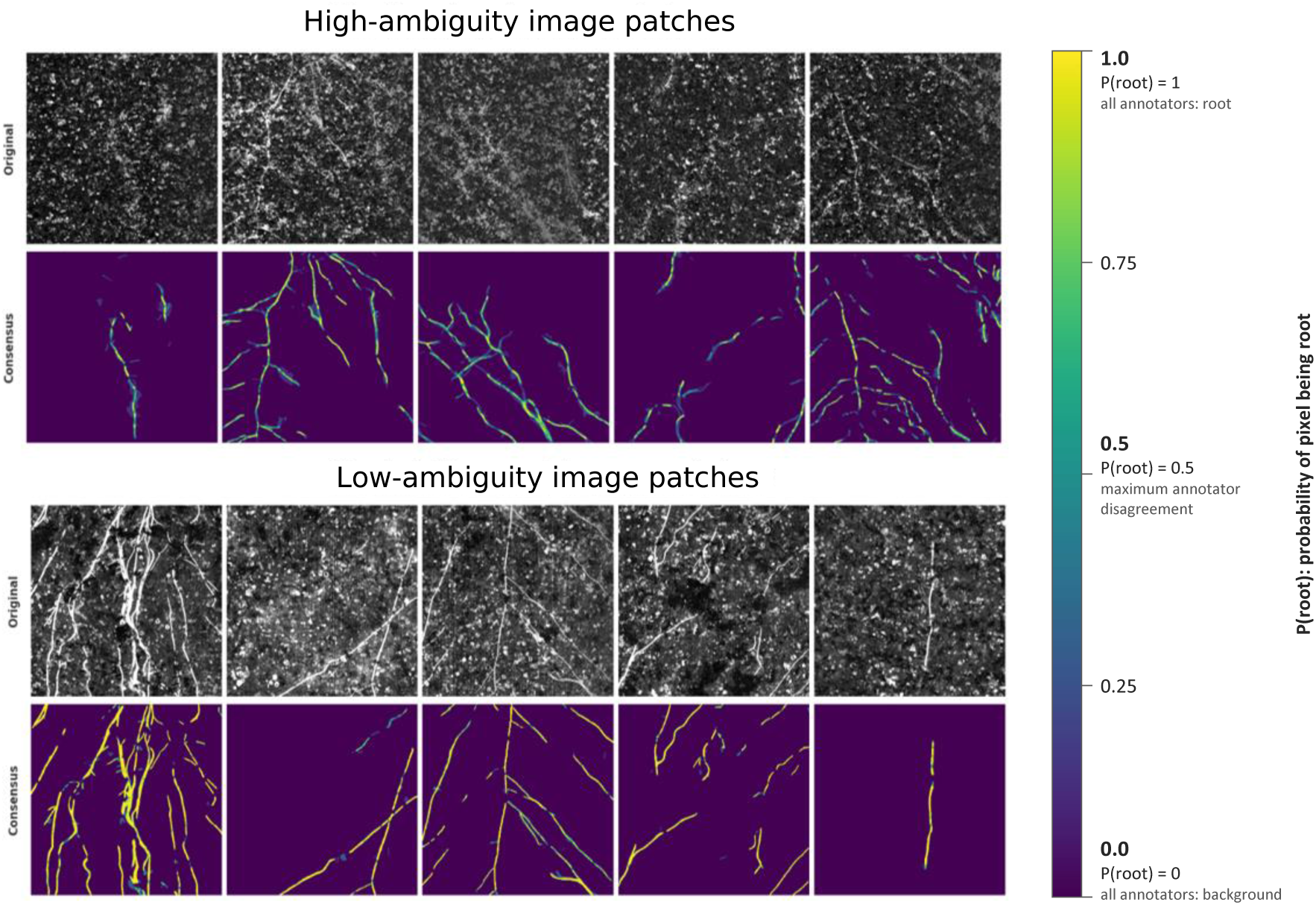
Probabilistic consensus maps for high-ambiguity and low-ambiguity image patches. Pixel values show the consensus probability *P* (*x*) (Equation 5; labelled *P* (root) in the figure), the proportion of the seven annotators labelling the pixel as root, from 0 (none) to 1 (all).

The corresponding Soft Dice scores are shown in Figure 9. The model results shown here correspond to models fine-tuned using 100% of the real training dataset. For the high-ambiguity patches, the average human annotator achieved a Soft Dice score of 0.678, compared with 0.603 for nnU-Net v2, 0.527 for YOLO26, and 0.506 for Mask2Former. Thus, all three segmentation models remained below the average human agreement on these patches. Scores for all training strategies are given in Supplementary Tables S5–S6.

**Figure 9:**
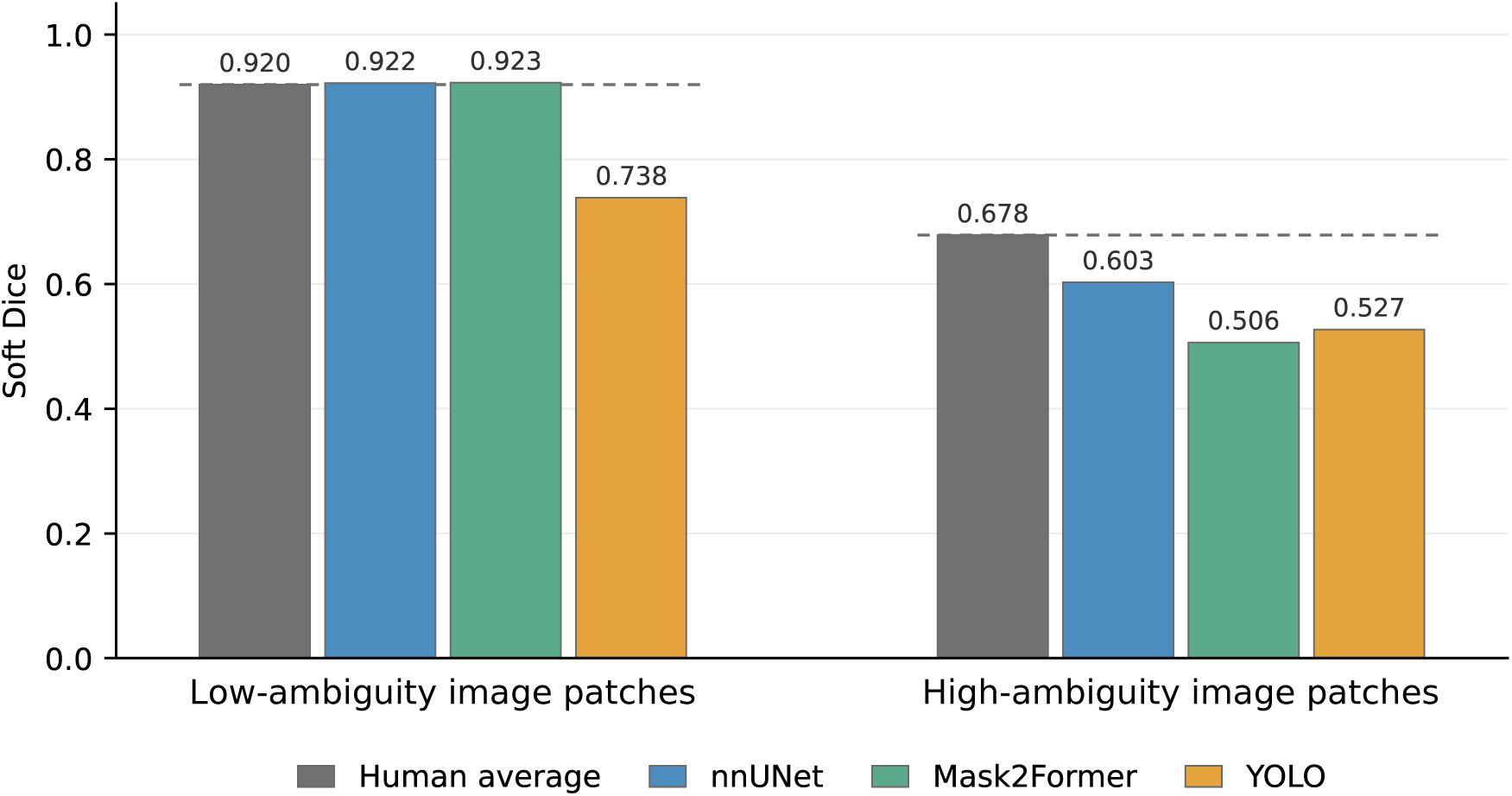
Soft Dice scores of human annotators and segmentation models. Scores are for low-ambiguity and high-ambiguity Arabidopsis patches. Models (nnU-Net v2, labelled nnUNet; Mask2Former; YOLO26, labelled YOLO) were fine-tuned on 100% of the real training data. Human and model scores use the leave-one-out consensus maps; model scores are averaged over all seven maps.

For the low-ambiguity image patches, substantially higher agreement was observed for both human annotators and segmentation models. The average human annotator achieved a Soft Dice score of 0.920, while nnU-Net v2 and Mask2Former achieved 0.922 and 0.923, respectively, slightly exceeding the average human score. YOLO26 achieved a lower Soft Dice score of 0.738. These results indicate that, when root structures are clearly distinguishable, some segmentation models can achieve performance comparable to or slightly above the average level of human agreement.

Overall, the comparison shows a clear dependence of model-human agreement on image ambiguity. On low-ambiguity patches, nnU-Net v2 and Mask2Former reached approximately human-level Soft Dice scores, whereas a substantial gap remained between all models and the human annotators on high-ambiguity patches containing ambiguous root boundaries, fine roots, and root hairs. The variation among human annotations on these high-ambiguity patches further indicates that segmentation performance in such cases is affected not only by model errors but also by uncertainty in the reference annotations. Detailed results for the different training strategies are provided in Supplementary Tables S5 and S6.

## 4 Discussion

Deep learning provides an opportunity to automate root segmentation in rhizotron images, but developing reliable models remains dependent on the availability of annotated training data. This study investigated the use of diffusion-generated synthetic rhizotron images to reduce this annotation requirement and support the development of transferable root segmentation models. The results show that synthetic data can serve as a complementary source of training information, supporting effective segmentation with substantially less manually labour intensive annotated real data.

### 4.1 Synthetic Pretraining Reduces Annotation Requirements

The results under the low-annotation settings show that synthetic pretraining can substantially reduce the amount of real data required to obtain effective root segmentation. Models pretrained on synthetic images generally outperformed models trained on the same limited amount of real data alone. This indicates that synthetic pretraining provides a useful initialisation, allowing models to learn general characteristics of root morphology before being adapted to the appearance of real rhizotron images.

This finding has practical significance for large-scale root phenotyping. Complete root annotation requires manually tracing potentially complex root systems and can become time-consuming even for a relatively small dataset, making extensive manual annotation increasingly impractical as the number of images grows. In our multi-annotator experiment, annotating the set of 10 high-ambiguity patches required approximately 1–3 h, even when using the semi-automated V7 Darwin platform based on SAM 3, whereas the set of 10 low-ambiguity patches required approximately 10 min–1.5 h. The high-ambiguity patches were not only more time-consuming to annotate but also showed greater disagreement among annotators, highlighting the difficulty of obtaining consistent reference annotations in visually ambiguous regions. At the lowest annotation level, the training set contained only 9 real patches, including just 2 patches with visible roots, and synthetic pretraining increased Dice from 0.573 to 0.623 for nnU-Net v2 and from 0.460 to 0.624 for Mask2Former. These results suggest that synthetic pretraining can be particularly valuable when real annotation is scarce and difficult to obtain, although the amount of annotation required for a given level of performance remains dependent on the segmentation model and characteristics of the target dataset.

The benefit was less pronounced when more real data was available. For nnU-Net v2, Dice increased from 0.786 with real-only training to 0.799 after synthetic pretraining, while clDice increased from 0.864 to 0.879. With sufficient real data, the model can learn the characteristics of the target domain directly, leaving less scope for additional information from synthetic pretraining. Once trained, the segmentation models can also be applied across large numbers of images using the same automated procedure, which can reduce the need for repeated manual tracing in large rhizotron datasets.

### 4.2 Quality and Characteristics of Synthetic Images

The synthetic-only experiments provide insight into the information captured by the generated images. Although performance varied between segmentation architectures, meaningful root segmentation was possible without exposure to real training images. In particular, YOLO26 achieved a Dice score of 0.596 when trained exclusively on synthetic images, while nnU-Net v2 achieved a Dice score of 0.416. The relatively strong performance of YOLO26 indicates that the generated dataset captured sufficient visual and structural information to support root segmentation in the absence of real training data. The difference between architectures also suggests that the extent to which synthetic information can be exploited depends partly on the segmentation model.

The synthetic-only results indicate that the generated images reproduce several characteristics relevant to the segmentation task, including root shape, branching structure, and visual differences between roots and surrounding soil. The further improvement obtained after fine-tuning with real images suggests that these representations provide a useful starting point for adapting the models to the target image domain.

Nevertheless, the difference between synthetic-only and real-data performance indicates that the generated images do not yet cover the full variability of real rhizotron images. Real images contain complex variation in root thickness and density, fine-root visibility, soil texture, illumination, occlusion, cracks, and root-soil contrast. The lower recall observed for synthetic-only models is consistent with this interpretation, as the models were able to identify plausible root structures but did not recover all structures present in real images. These findings suggest that improving the diversity of synthetic data, including a broader range of root architectures, root densities, soil textures, illumination conditions, and occlusion patterns, may be more important than simply increasing its size.

### 4.3 Synthetic Pretraining and Cross-Species Generalisation

The models trained primarily on Arabidopsis data were able to segment several previously unseen crop species without additional fine-tuning. This suggests that the learned representations capture structural characteristics of roots that are shared across species rather than being restricted to species-specific appearance.

The benefit of synthetic pretraining was particularly evident when the amount of real training data was limited. One possible explanation is that the synthetic dataset exposes the models to a wider range of root configurations than would be available from a small real training set. This may encourage the models to learn general features related to root continuity, tubular structure, and branching, which can subsequently be transferred to other species.

The extent of transfer varied between crop species. Differences in root thickness, branching architecture, density, soil composition, and imaging conditions can introduce domain shifts that affect segmentation performance. The current results should therefore be interpreted as evidence of promising cross-species transfer rather than complete species-independent generalisation. Nevertheless, obtaining useful segmentation of unseen crops without creating a large species-specific annotated dataset is practically relevant, and suggests that synthetic pretraining could facilitate the adaptation of root segmentation models to new crops where only limited annotated data is available.

### 4.4 Human Annotation Variability and Evaluation Uncertainty

The evaluation results also highlight an important characteristic of the root segmentation task: manual annotation is itself subject to substantial uncertainty. Differences among the seven annotators were particularly apparent in images containing fine roots, weak root-soil contrast, blurred boundaries, and visually ambiguous structures. Consequently, disagreement between a model prediction and an individual annotation does not necessarily represent an unequivocal segmentation error. A conventional binary ground truth assumes that each pixel can be assigned reliably to either root or soil, whereas this assumption is not always appropriate for visually ambiguous rhizotron images. A model prediction that differs from one annotator may still be consistent with the interpretation of another.

To account for this uncertainty, we derived a probabilistic consensus map from the seven independent annotations and used it to calculate Soft Dice. This approach assigns greater weight to regions where annotators agree and lower weight to regions with greater disagreement. Models approached the performance of individual annotators for low-ambiguity patches containing clearly distinguishable root structures, while a larger difference remained for high-ambiguity patches. Segmentation performance should therefore be interpreted relative to the level of human agreement rather than against an assumption of error-free ground truth.

### 4.5 Limitations and Steps Toward Application

Several limitations should be considered when interpreting these results. The real training data were derived from only two annotated full rhizotron images, annotated by a single primary annotator, and all images were acquired with one rhizotron platform. The absolute scores may therefore differ for other platforms or annotators, but the comparison between training strategies, on which our conclusions are based, is less affected because all strategies used the same data and reference annotations. The synthetic root structures were recombined from Arabidopsis root masks annotated on agar plates, which limits the range of root architectures represented; this probably contributes to the gap between synthetic-only and real-data models and may limit transfer to crops with very different root systems. Synthetic pretraining did not benefit YOLO26 in most settings, so the gains reported here depend on the model architecture. Finally, the human-level comparison was based on 20 image patches from seven annotators and should be regarded as indicative; within this set, models remained below average human agreement on high-ambiguity patches.

Most importantly, we evaluated pixel-level and topology-aware segmentation accuracy but did not assess how differences in model predictions propagate to downstream root traits. Two masks may differ substantially at the pixel level while producing similar estimates of root length, whereas small differences in particular regions could affect other measurements disproportionately. For the findings to be applied in phenotyping studies, the relationship between segmentation accuracy and the accuracy of derived root traits should be validated against reference measurements, and the approach should be tested quantitatively across rhizotron platforms, soil conditions, and imaging protocols. Preliminary experiments using different rhizotron systems showed promising transferability (Supplementary Figs. S3–S4).

### 4.6 Future Directions

The gap between synthetic-only and real-data performance shows that the realism of the generated images still needs to be improved. Several current issues can be addressed directly. First, the root structures are recombined from Arabidopsis root masks annotated on agar plates, so their architecture, root thickness and branching do not fully represent soil-grown roots or other crops. Conditioning masks derived from soil-grown roots of several species, or generated procedurally, for example with L-system root models as used by Hann et al. [13], could broaden the range of architectures while reducing the effort needed to prepare them. Second, the Canny edge map passed to ControlNet constrains the root outline but not root diameter or contrast along the root, so additional conditioning signals, such as root-width or distance maps, could give more control over root appearance. Third, the Root LoRA and the soil patches were derived from one rhizotron platform and one species, so training them on images from several platforms, soils and illumination conditions would increase the variability that the generated images cover. Finally, realism was assessed here only through downstream segmentation performance; complementary measures, such as image-distribution metrics or blinded comparisons by experts, would help to identify which characteristics of real images are still missing.

An important issue for application is root occlusion. In rhizotron images, roots are often partly hidden behind soil particles or grow away from the transparent window, so a root that is continuous in reality appears as separate visible segments. The current pipeline generates roots along the entire root-mask patch, so the synthetic images assume that every part of the root is visible, and the models were trained and evaluated only on visible root pixels. For trait measurements such as total root length or root counts, however, the hidden parts also need to be inferred. The synthetic approach is well suited to this problem, because the complete root architecture is known for every generated image. Parts of the roots could be deliberately covered with inpainted soil while the complete root-mask patch is retained as the label, producing paired training data for models that predict both visible and occluded roots. Topology-aware metrics such as clDice could then be used to assess whether the predicted root systems remain connected across occluded regions.

Synthetic pretraining could also be combined with active learning and uncertainty-aware training. Active learning could identify the images or regions where additional annotation would be most informative, while probabilistic labels or multi-annotator supervision could allow models to account for ambiguity in root-soil boundaries.

## 5 Conclusions

We developed a diffusion-based framework for generating synthetic rhizotron images, together with a Soft Dice metric that evaluates segmentation against probabilistic consensus maps from multiple annotators, and found that (i) synthetic images contained sufficient information to support meaningful root segmentation, although performance remained below that of models trained on real annotated data, indicating that the generated images do not yet capture the full variability of real rhizotron images and cannot replace real annotations; (ii) synthetic pretraining improved segmentation under annotation-limited conditions, whereas the benefit was small when the full real training set was available and depended on the segmentation architecture; (iii) synthetic pretraining improved generalisation to unseen crop species when real data were limited, although the gain varied between species; and (iv) models matched average human agreement on low-ambiguity patches but remained below it on high-ambiguity patches, where human-level evaluation also showed substantial inter-annotator variability; the consensus maps and Soft Dice take this variability into account and allow model performance to be interpreted relative to human agreement. Combining synthetic pretraining with a small amount of real annotated data can therefore reduce the manual annotation burden, which is currently a key bottleneck for root segmentation in high-throughput rhizotron phenotyping, and help models trained on a few annotated images to be applied automatically to large image series. Further work should improve the realism of the generated images, including occluded roots, test the approach across rhizotron platforms, and validate its effect on downstream root traits.

## Declaration of generative AI and AI-assisted technologies in the manuscript preparation process

During the preparation of this work the authors used ChatGPT (OpenAI) to assist with English-language editing, coding, and figure improvement, and Claude (Anthropic) to assist with English-language editing, coding, and organising and documenting the code repository. After using these tools, the authors reviewed and edited the content as needed and take full responsibility for the content of the published article. The synthetic rhizotron images shown in Figures 1, 2 and 4 were generated with Stable Diffusion 1.5 as described in Materials and Methods; they are synthetic training data and not experimental observations.

## Data availability

The code for root-mask composition, Root LoRA training, synthetic image generation, segmentation model training and evaluation (including the Soft Dice metric) will be made publicly available on GitHub upon journal publication. The synthetic rhizotron dataset (3,758 images with their root masks), the Root LoRA weights and the trained segmentation models will be made publicly available on Zenodo upon journal publication. The real rhizotron images used in this study were acquired in collaboration with an industrial partner and are subject to a confidentiality agreement; they therefore cannot be made publicly available.

## Acknowledgments

We acknowledge the Netherlands Plant Eco-phenotyping Centre (NPEC) for providing the rhizotron imaging platform and technical support. We gratefully acknowledge all volunteers (Iva Andrasevic, Thijs Stegmann, Anupama Balagopal, Hazel van Waijjen, Filip Toncic, and Chenxin Luo) who contributed to the manual annotation of the rhizotron images. Their efforts were essential for constructing the reference datasets and conducting the inter-annotator evaluation presented in this study.

## CRediT authorship contribution statement

**Fengjiao Lu:** Conceptualization, Methodology, Software, Validation, Formal analysis, Investigation, Data curation, Visualization, Writing – original draft, Writing – review & editing. **Gert Kootstra:** Conceptualization, Supervision, Writing – review & editing. **Zane K. J. Hartley:** Conceptualization, Writing – review & editing. **Rick van de Zedde:** Conceptualization, Supervision. **Michael Pound:** Conceptualization, Supervision, Writing – review & editing. **Viola Willemsen:** Conceptualization, Supervision, Writing – review & editing.

