## Supplementary files for "Diffusion-Based Synthetic Pretraining for Root Segmentation in Rhizotron Images"

#### Detailed segmentation performance

Tables S1, S2 and S3 present the complete segmentation results on the Arabidopsis test set for nnU-Net v2, YOLO26, and Mask2Former under all training strategies and annotation levels. Synthetic-only, real-only and PT+FT (synthetic pretraining followed by fine-tuning on real data) are the training strategies defined in the main text; annotation levels are proportions of annotated root pixels in the real training dataset. In addition to Dice and clDice, the tables report Precision and Recall, providing a more comprehensive evaluation of segmentation performance.

Overall, synthetic pretraining gave a clear benefit only at the 10% annotation level, for nnU-Net v2 and Mask2Former. At 25–75% and 100% the differences between real-only and PT+FT models were small and not consistent in sign. The Precision and Recall values further indicate that nnU-Net v2 and Mask2Former trained exclusively on synthetic data achieved high precision (0.862 and 0.790) but low recall (0.278 and 0.251), i.e. they detected only part of the root pixels present in real images, consistent with a remaining domain gap between synthetic and real rhizotron images. YOLO26 was more balanced (precision 0.722, recall 0.533).

Table S1: Detailed segmentation performance of nnU-Net v2 on the Arabidopsis test set under different training strategies and annotation levels. Synthetic-only, trained on synthetic data only; real-only, trained on the real-data subset only; PT+FT, pretrained on synthetic data and fine-tuned on the same real-data subset.

| **Training strategy** | **Annotation level** | **Dice** | **clDice** | **Precision** | **Recall** |
|---|---|---|---|---|---|
| Synthetic-only | 0% (no real data) | 0.416 | 0.672 | 0.862 | 0.278 |
| Real-only | 100% (2 images) | 0.786 | 0.864 | 0.847 | 0.757 |
| | 75% | 0.791 | 0.868 | 0.839 | 0.767 |
| | 50% | 0.782 | 0.858 | 0.850 | 0.746 |
| | 25% | 0.760 | 0.837 | 0.842 | 0.720 |
| | 10% | 0.573 | 0.643 | 0.702 | 0.511 |
| PT+FT | 100% | 0.799 | 0.879 | 0.843 | 0.778 |
| | 75% | 0.796 | 0.875 | 0.837 | 0.778 |
| | 50% | 0.772 | 0.855 | 0.858 | 0.725 |
| | 25% | 0.742 | 0.818 | 0.827 | 0.704 |
| | 10% | 0.623 | 0.703 | 0.831 | 0.530 |

Table S2: Detailed segmentation performance of YOLO26 on the Arabidopsis test set under different training strategies and annotation levels. Synthetic-only, trained on synthetic data only; real-only, trained on the real-data subset only; PT+FT, pretrained on synthetic data and fine-tuned on the same real-data subset.

| Training strategy | Annotation level | Dice | clDice | Precision | Recall |
|---|---|---|---|---|---|
| Synthetic-only | 0% (no real data) | 0.596 | 0.689 | 0.722 | 0.533 |
| Real-only | 100% (2 images) | 0.671 | 0.838 | 0.576 | 0.816 |
| | 75% | 0.623 | 0.779 | 0.568 | 0.722 |
| | 50% | 0.634 | 0.809 | 0.551 | 0.769 |
| | 25% | 0.638 | 0.789 | 0.578 | 0.735 |
| | 10% | 0.659 | 0.781 | 0.636 | 0.713 |
| PT+FT | 100% | 0.662 | 0.836 | 0.562 | 0.820 |
| | 75% | 0.636 | 0.794 | 0.558 | 0.766 |
| | 50% | 0.611 | 0.763 | 0.511 | 0.780 |
| | 25% | 0.614 | 0.776 | 0.515 | 0.770 |
| | 10% | 0.670 | 0.806 | 0.622 | 0.757 |

Table S3: Detailed segmentation performance of Mask2Former on the Arabidopsis test set under different training strategies and annotation levels. Synthetic-only, trained on synthetic data only; real-only, trained on the real-data subset only; PT+FT, pretrained on synthetic data and fine-tuned on the same real-data subset.

| Training strategy | Annotation level | Dice | clDice | Precision | Recall |
|---|---|---|---|---|---|
| Synthetic-only | 0% (no real data) | 0.375 | 0.605 | 0.790 | 0.251 |
| Real-only | 100% (2 images) | 0.748 | 0.817 | 0.836 | 0.700 |
| | 75% | 0.754 | 0.830 | 0.845 | 0.699 |
| | 50% | 0.728 | 0.812 | 0.869 | 0.649 |
| | 25% | 0.710 | 0.789 | 0.820 | 0.647 |
| | 10% | 0.460 | 0.527 | 0.671 | 0.364 |
| PT+FT | 100% | 0.747 | 0.803 | 0.797 | 0.726 |
| | 75% | 0.740 | 0.797 | 0.802 | 0.712 |
| | 50% | 0.747 | 0.825 | 0.846 | 0.691 |
| | 25% | 0.654 | 0.704 | 0.622 | 0.708 |
| | 10% | 0.624 | 0.691 | 0.689 | 0.597 |

### Generalisation to different crop species

To provide a more detailed view of the cross-species evaluation, Figures S1 and S2 present the segmentation performance for each species in the cross-species test set (Arabidopsis and eight crop species). All models were trained only on Arabidopsis data (real Arabidopsis images, with or without synthetic pretraining) and evaluated directly on the cross-species test set without additional fine-tuning or domain adaptation.

Figure S1 compares the synthetic-only model with the real-only and PT+FT models trained using the full (100%) real Arabidopsis training dataset, whereas Figure S2 compares the real-only and PT+FT models when only 10% of the real training data were available. The figures report both Dice and clDice scores for each crop species and segmentation model.

Average Dice scores across all nine species are summarised in Table S4.

Table S4: Average Dice scores by training strategy and annotation level on the cross-species test set (averaged over Arabidopsis and the eight crop species). Real-only, trained using real images only; PT+FT, pretrained on synthetic data and fine-tuned on real data. Bold indicates the higher score within each model and annotation level.

| Model | 10% real data | | 100% real data | |
|---|---|---|---|---|
| | Real-only | PT+FT | Real-only | PT+FT |
| nnU-Net v2 | 0.515 | **0.536** | **0.676** | 0.667 |
| YOLO26 | 0.599 | **0.626** | 0.648 | **0.668** |
| Mask2Former | 0.334 | **0.441** | **0.604** | 0.602 |

Overall, synthetic pretraining provided the largest improvements under the 10% annotation setting, particularly for Mask2Former, while the performance gains became smaller when the full real training dataset was available. Performance varied across crop species, reflecting differences in root morphology, soil appearance, and imaging conditions, and synthetic pretraining did not improve every species for every model (e.g., decreases for field bean and tomato with nnU-Net v2 and for wheat with YOLO26 under the 10% setting).

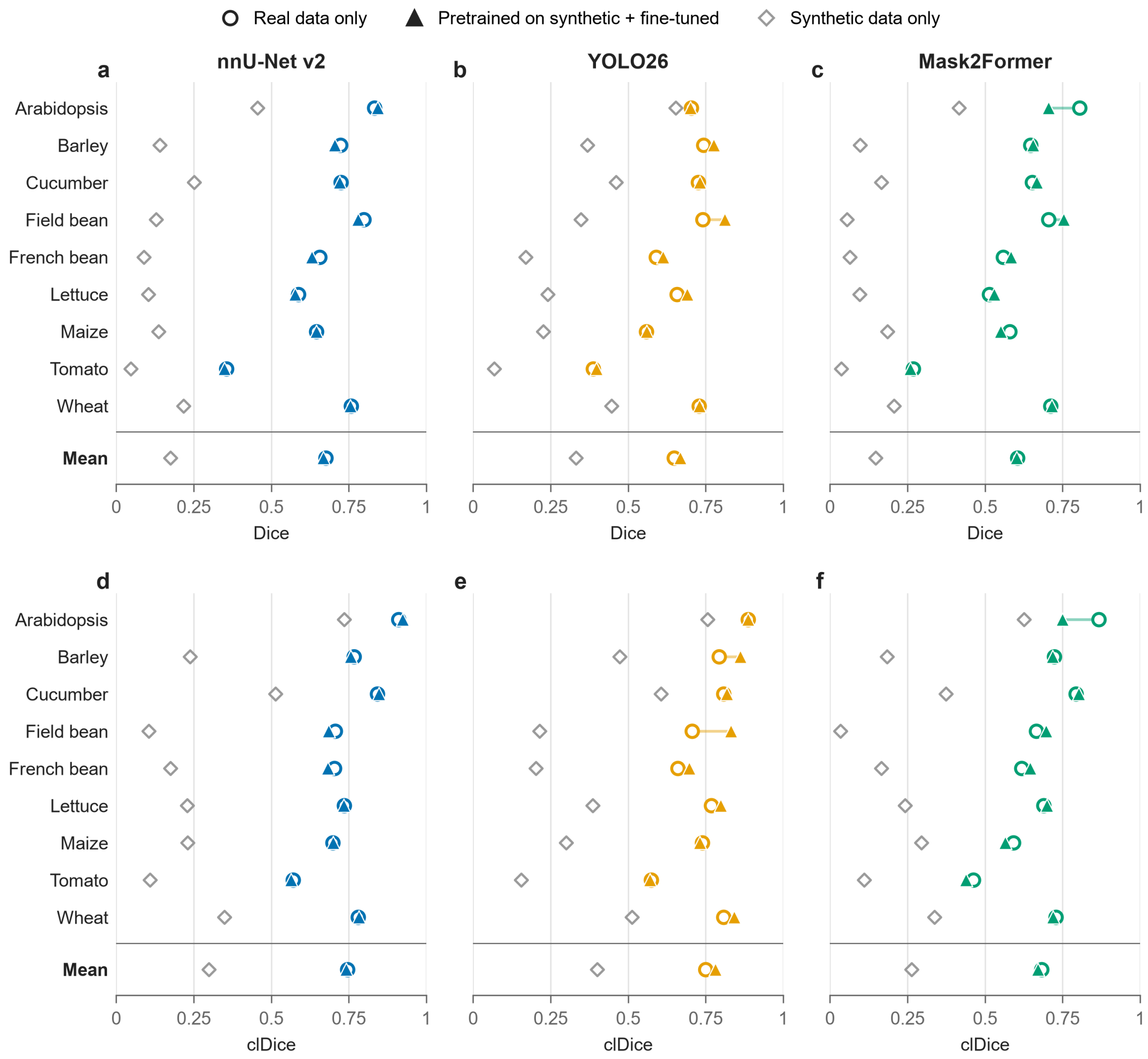


Figure S1: Segmentation performance per species in the cross-species test set with 100% of the real Arabidopsis training dataset. (a–c) Dice and (d–f) clDice for nnU-Net v2, YOLO26 and Mask2Former. Open circles: real-only models; filled triangles: PT+FT models; the line between them shows the effect of synthetic pretraining. Grey diamonds: synthetic-only models (no real data). "Mean" is the average over the nine species (Supplementary Table S4). Barley is cultivar Planet.

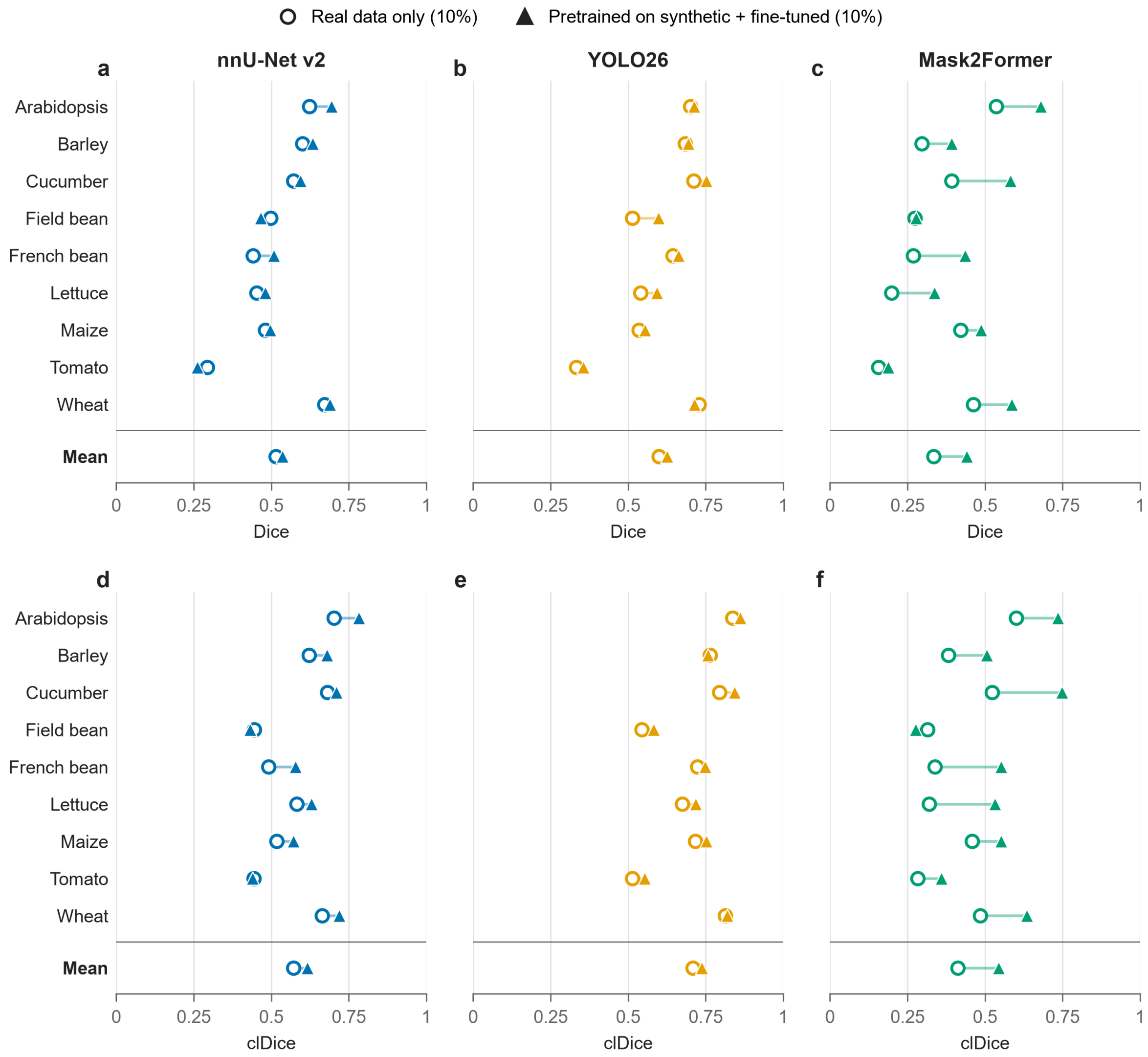


Figure S2: Segmentation performance per species in the cross-species test set with 10% of the real Arabidopsis training dataset. (a–c) Dice and (d–f) clDice for nnU-Net v2, YOLO26 and Mask2Former. Open circles: real-only models; filled triangles: PT+FT models; the line between them shows the effect of synthetic pretraining. "Mean" is the average over the nine species (Supplementary Table S4). Barley is cultivar Planet.

### Detailed human annotator performance

Tables S5 and S6 report Soft Dice, Precision, Recall and Soft clDice against the probabilistic consensus maps for the 10 low-ambiguity and 10 high-ambiguity Arabidopsis patches (Experiment 4). Synthetic-only denotes training on synthetic data only; Real-only ($N$%) denotes training on the $N$% annotation level of the real data only; PT+FT ($N$%) denotes synthetic pretraining followed by fine-tuning on the same $N$% subset. "Human average" is the mean over annotators, each scored against a leave-one-out consensus of the remaining six annotators. Best values per column are in bold.

All metrics use the uncertainty weight $w(x) = (1-w_{\min})\, 2|P(x)-0.5|+w_{\min}$ with $w_{\min} = 0.1$, where $P(x)$ is the consensus probability and $V_P(x) \in \{0, 1\}$ the evaluated mask, as in the main text. Soft Precision and Soft Recall are computed from the weighted true positives $\sum wPV_P$, false positives $\sum w(1-P)V_P$ and false negatives $\sum wP(1-V_P)$. Soft clDice is a weighted version of clDice: with $S_L$ the skeleton of the consensus map binarised at $P \geq 0.5$ and $S_P$ the skeleton of $V_P$,

$$T_{\text{sens}} = \frac{\sum wPS_LV_P}{\sum wPS_L}, \qquad T_{\text{prec}} = \frac{\sum wPS_P}{\sum wS_P}, \qquad \text{Soft clDice} = \frac{2\,T_{\text{prec}}T_{\text{sens}}}{T_{\text{prec}} + T_{\text{sens}}},$$

so $T_{\text{sens}}$ is the weighted fraction of the consensus skeleton covered by the mask and $T_{\text{prec}}$ the weighted fraction of the mask skeleton that lies on consensus roots. Each model mask is scored against all seven leave-one-out consensus maps and the scores are averaged; all metrics are computed per patch and averaged over the 10 patches.

Table S5: Detailed performance of human annotators and segmentation models on the 10 low-ambiguity Arabidopsis image patches.

| **Method** | **Soft Dice** | **Soft Precision** | **Soft Recall** | **Soft clDice** |
|---|---|---|---|---|
| Mask2Former - Synthetic-only | 0.5280 | **0.9708** | 0.3740 | 0.8129 |
| nnU-Net v2 - Synthetic-only | 0.5355 | 0.8791 | 0.3977 | 0.7775 |
| YOLO26 - Real-only (10%) | 0.6669 | 0.5871 | 0.7808 | 0.7728 |
| YOLO26 - Synthetic-only | 0.6714 | 0.7486 | 0.6320 | 0.7098 |
| YOLO26 - Real-only (100%) | 0.7213 | 0.5968 | 0.9210 | 0.8964 |
| YOLO26 - PT+FT (100%) | 0.7384 | 0.5979 | **0.9677** | 0.9080 |
| Mask2Former - Real-only (10%) | 0.7498 | 0.9451 | 0.6477 | 0.8120 |
| YOLO26 - PT+FT (10%) | 0.7596 | 0.6534 | 0.9179 | 0.9064 |
| Mask2Former - PT+FT (10%) | 0.8590 | 0.9363 | 0.8049 | 0.9287 |
| nnU-Net v2 - PT+FT (10%) | 0.8653 | 0.9070 | 0.8420 | 0.9273 |
| nnU-Net v2 - Real-only (10%) | 0.8701 | 0.9170 | 0.8370 | 0.9297 |
| Mask2Former - Real-only (100%) | 0.9155 | 0.8990 | 0.9340 | 0.9637 |
| nnU-Net v2 - Real-only (100%) | 0.9198 | 0.8943 | 0.9474 | 0.9685 |
| nnU-Net v2 - PT+FT (100%) | 0.9222 | 0.8945 | 0.9521 | **0.9720** |
| Mask2Former - PT+FT (100%) | **0.9229** | 0.9059 | 0.9416 | 0.9687 |
| Human average | 0.9198 | 0.9237 | 0.9204 | 0.9704 |

Table S6: Detailed performance of human annotators and segmentation models on the 10 high-ambiguity Arabidopsis image patches.

| **Method** | **Soft Dice** | **Soft Precision** | **Soft Recall** | **Soft clDice** |
|---|---|---|---|---|
| Mask2Former - Synthetic-only | 0.2491 | 0.8010 | 0.1594 | 0.3936 |
| Mask2Former - Real-only (10%) | 0.2648 | **0.9110** | 0.1755 | 0.3293 |
| YOLO26 - Synthetic-only | 0.2836 | 0.4690 | 0.2360 | 0.3153 |
| nnU-Net v2 - Synthetic-only | 0.3180 | 0.8524 | 0.2168 | 0.4912 |
| nnU-Net v2 - PT+FT (10%) | 0.3717 | 0.9049 | 0.2595 | 0.4575 |
| nnU-Net v2 - Real-only (10%) | 0.4034 | 0.8743 | 0.2795 | 0.5012 |
| Mask2Former - PT+FT (10%) | 0.4121 | 0.8110 | 0.3087 | 0.4978 |
| YOLO26 - Real-only (10%) | 0.4558 | 0.6090 | 0.4188 | 0.5451 |
| YOLO26 - PT+FT (10%) | 0.4563 | 0.6072 | 0.4334 | 0.5574 |
| Mask2Former - PT+FT (100%) | 0.5062 | 0.7765 | 0.4187 | 0.5972 |
| YOLO26 - Real-only (100%) | 0.5169 | 0.5422 | 0.5449 | 0.6699 |
| Mask2Former - Real-only (100%) | 0.5238 | 0.8638 | 0.4200 | 0.6276 |
| YOLO26 - PT+FT (100%) | 0.5269 | 0.5129 | 0.5762 | 0.6993 |
| nnU-Net v2 - Real-only (100%) | 0.5692 | 0.8663 | 0.4676 | 0.6843 |
| nnU-Net v2 - PT+FT (100%) | 0.6028 | 0.8616 | 0.5055 | 0.7286 |
| Human average | **0.6784** | 0.6852 | **0.6948** | **0.8341** |

### Generalisation to different rhizotron systems

To provide a preliminary assessment of model transferability, we evaluated the synthetic-pretrained model on images acquired from two independent rhizotron systems that were not included in the training data. The predictions in Figures S3 and S4 were produced with nnU-Net v2. The model was pretrained on synthetic Arabidopsis images and fine-tuned on the full real Arabidopsis training set (100% setting; 108 patches from two annotated rhizotron images) before being directly applied to the new datasets without additional fine-tuning.

The first dataset consisted of barley rhizotron images acquired using a different rhizotron system (Figure S3). The second dataset consisted of Arabidopsis images acquired using another independent rhizotron system (Figure S4). Although no quantitative evaluation was performed, the representative predictions demonstrate that the model is able to identify the major root structures under different imaging conditions and across different rhizotron platforms. These preliminary results suggest that the proposed framework has the potential to generalise beyond the imaging system used for training.

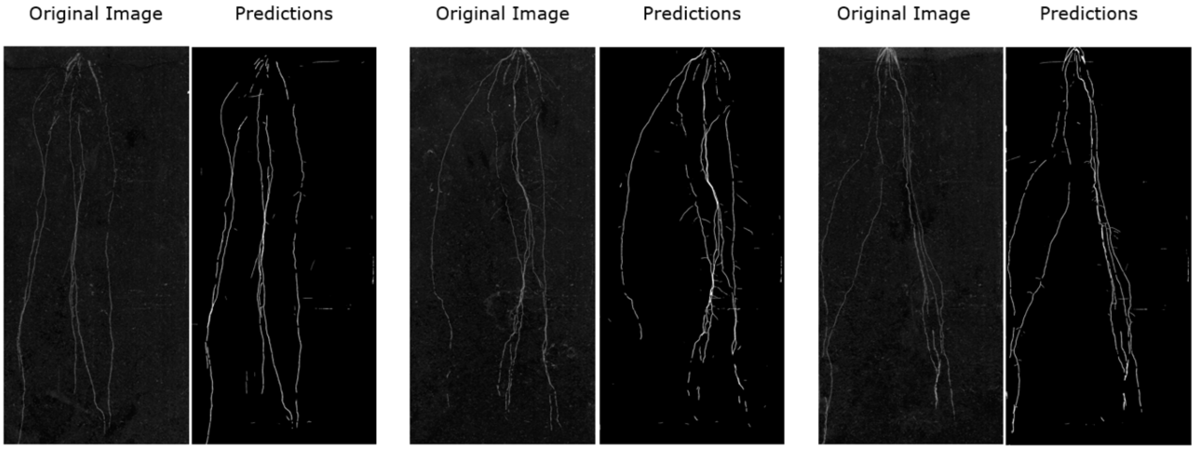


Figure S3: Preliminary predictions on barley rhizotron images acquired using an independent rhizotron system. Predictions were produced with the nnU-Net v2 PT+FT model (100% annotation level) without additional fine-tuning.

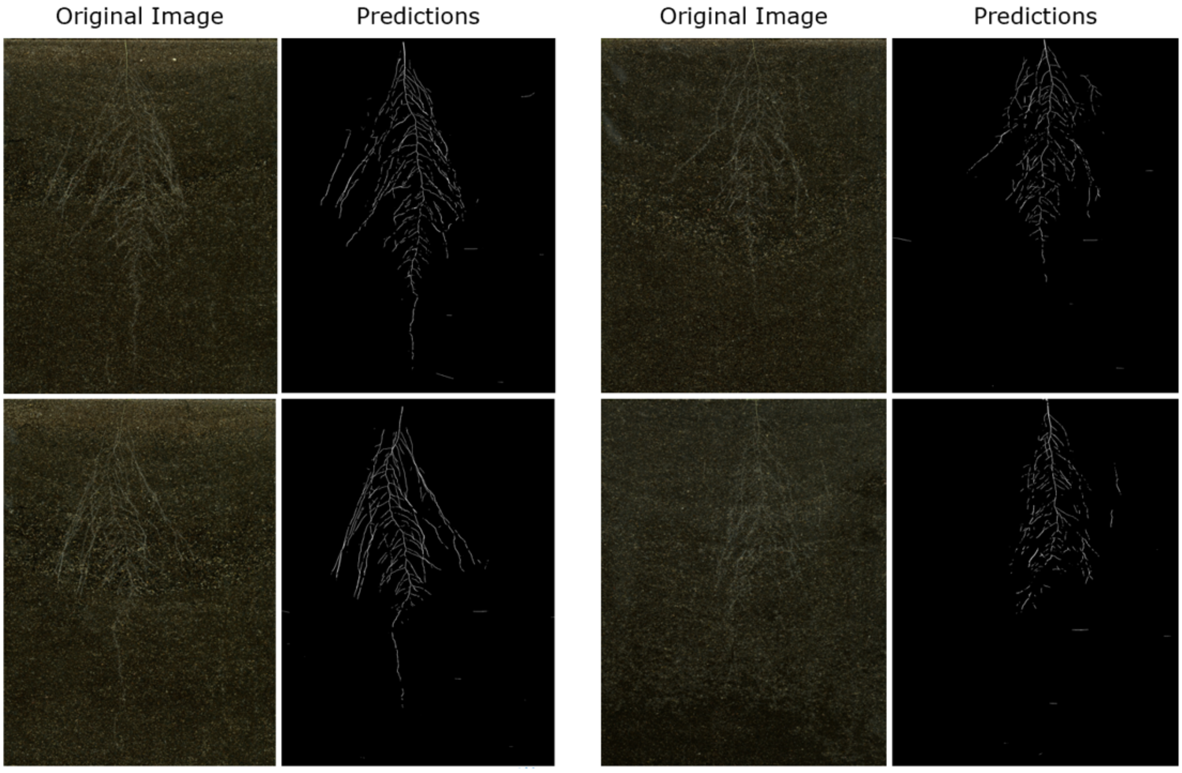


Figure S4: Preliminary predictions on Arabidopsis rhizotron images acquired using a second independent rhizotron system with a different type of soil. Predictions were produced with the nnU-Net v2 PT+FT model (100% annotation level) without additional fine-tuning.
